# Nimodipine reduces microglial activation *in vitro* as evidenced by morphological phenotype, phagocytic activity and next generation RNA sequencing

**DOI:** 10.1101/2024.11.11.623031

**Authors:** István Pesti, Valentin Varga, Erda Qorri, Rita Frank, Diana Kata, Krisztián Vinga, Péter Archibald Szarvas, Ákos Menyhárt, Károly Gulya, Ferenc Bari, Eszter Farkas

**Author notes:** Correspondence Eszter Farkas, D.Sc., HCEMM-USZ Cerebral Blood Flow and Metabolism Research Group Department of Cell Biology and Molecular Medicine, Szent-Györgyi Albert Medical School Faculty of Science and Informatics University of Szeged, Somogyi u. 4 H-6720 Szeged Hungary.

## Abstract

**Background:** Nimodipine, an L-type voltage-gated calcium channel blocker, achieves vasorelaxation by suppressing Ca^2+^-dependent activation of cerebrovascular smooth muscle cells and is used to prevent delayed ischemic deficit following subarachnoid hemorrhage. Our preclinical drug repurposing studies raised the possibility that nimodipine may attenuate the pro-inflammatory shift in microglial function in response to brain injury. We analyzed the effects of nimodipine on activated microglia at the level of morphological and functional phenotypes, as well as their transcriptomic profile.

**Methods:** Live brain slice preparations from C57BL/6 mice and primary microglia cultures from the cortex of neonatal Sprague Dawley rats were used. Brain slices were subjected to ischemia, and microglial cultures were activated with lipopolysaccharide (LPS; 20 ng/ml). Both preparations were treated with nimodipine (5-10-20 μM). The degree of arborization was evaluated in Iba1-stained microglia and expressed as a transformation index (TI). Phagocytic activity of cultured microglia was visualized using fluorescent microbeads. TNFα levels in the cultures were measured with ELISA. Total RNA was isolated from microglia and processed for next generation RNA sequencing to determine differentially expressed genes.

**Results:** Nimodipine suppressed the ameboid morphological transformation and increased phagocytosis triggered by ischemia in brain slices and LPS in microglia cultures. At the transcriptional level, LPS resulted in a pro-inflammatory microglial phenotype, affecting the expression of cytokines, the complement system and phagocytosis-related genes. Focusing on the role of calcium in microglial activation, LPS increased RNA transcription of ionotropic purinergic and some TRP channels but decreased the expression of voltage- and ligand-gated calcium channels. In the endoplasmic reticulum, LPS downregulated gene expression of Ryr and IP3 receptors and increased transcription of the SERCA calcium pump gene. Nimodipine co-administered with LPS altered the expression of 110 genes in the opposite direction to LPS activation, of which at least 20 were associated with microglial immune response, 7 with cell adhesion and 2 with autophagy regulation.

**Conclusion:** The effect of nimodipine goes beyond cerebral vasorelaxation. Nimodipine attenuates microglial activation by modulating Ca^2+^-dependent gene expression involved in intracellular signaling cascades to drive microglial immune responses. Consideration should be given to expanding the medical field of indication of nimodipine.

## Background

Nimodipine is a 1,4-dihydropyridine L-type voltage-gated calcium channel (LVGCC) blocker (Tang et al., 2016). The widely accepted primary targets of nimodipine are cerebrovascular smooth muscle cells (Kazda and Towart, 1982; Freedman and Waters, 1987). Nimodipine achieves vasorelaxation by inhibiting Ca^2+^ influx through LVGCCs, thereby reducing Ca^2+^-dependent activation of smooth muscle cell contractile activity. Nimodipine has clinical therapeutic value in the prevention of delayed ischemic deficit following subarachnoid hemorrhage (Allen et al., 1983; Pickard et al., 1989; Feigin et al., 1998). In addition, recent preclinical investigations have revisited the possibility that nimodipine may provide therapeutic benefit in selected patients with ischemic stroke or migraine with aura (Carlson et al., 2020), and have experimentally shown that a common pathomechanism known as spreading depolarization (SD) can be partially inhibited by nimodipine (Dietz et al., 2008; Menyhárt et al., 2018; Szabó et al., 2019; Frank et al., 2024; Menyhárt et al., 2024).

In addition to vascular smooth muscle cells, LVGCCs are expressed in various cell types of the nervous tissue, including microglia (Colton et al., 1994; Espinosa-Parrilla et al., 2015). Microglia are the resident immune cells in the central nervous system, which become rapidly activated by exogenous pathogens or endogenous danger signals upon brain injury. Activated microglia can assume functional states that are implicated in neuroinflammation and neurotoxicity. Importantly, microglial activation has been associated with altered intracellular Ca^2+^ homeostasis (Hopp et al., 2020). *In vivo* two-photon imaging identified distinct microglial Ca^2+^ transients in response to acute neuronal injury (Eichhoff et al., 2011) and prominent microglial Ca^2+^ waves associated with SD in experimental ischemic stroke (Liu et al., 2021). Exposure of cultured microglia to the bacterial lipopolysaccharide (LPS) revealed a sustained Ca^2+^ load of the cells, which blunted receptor-mediated Ca^2+^ transients (Hoffmann et al., 2003). At least part of the Ca^2+^ influx upon microglial activation must be mediated by LVGCCs. For example, various LVGCC blockers reduced the intracellular Ca^2+^ load in microglia activated with prion protein or beta-amyloid fragments (Silei et al., 1999), and the addition of nimodipine attenuated RANTES-induced Ca^2+^ accumulation in cultured microglia (Hegg et al., 2000). Further, a Ca^2+^ current via LVGCCs sensitive to nifedipine (a dihydropyridine calcium channel blocker similar to nimodipine) was associated with superoxide production by microglia (Colton et al., 1994). Finally, nimodipine administered to LPS-activated microglial cultures inhibited the production of the pro-inflammatory tumor necrosis factor-α (TNF-α), interleukin-1β (IL-1β) and prostaglandin E2 (PGE2) (Li et al., 2009). The effect of nimodipine on microglial LVGCCs is realistic because microglia express the Cav1.2 and Cav1.3 isoforms of the pore-forming α subunit of the channel (Espinosa-Parrilla et al., 2015; Hopp et al., 2020), for which nimodipine affinity has been described (Xu and Lipscombe, 2001; Huang et al., 2013). It remains to be determined, which intracellular signaling cascades link the pro-inflammatory shift in microglial phenotype to intracellular Ca^2+^ oscillations.

We have recently found that nimodipine given to the bath of metabolically challenged live brain slice preparations undergoing SD shifts the morphological phenotype of activated microglia to a more quiescent state, as evidenced by richer microglial ramifications (Frank et al., 2024). However, the model system does not allow to separate direct nimodipine effects on microglia from indirect actions (e.g. nimodipine altering neuronal excitation to which microglia respond), because LVGCCs are ubiquitously expressed in several cell types in the brain tissue, including neurons, astrocytes, and oligodendrocytes (Hopp et al., 2020). Primary microglial cultures have been successfully used to study microglial activation and the effect of different pharmacological agents on microglial states selectively (Pesti et al., 2024). Further, transcriptomic analysis of cultured microglia activated with LPS revealed the induction of pro-inflammatory genes, interferon-induced genes, and immune activation genes (Pulido-Salgado et al., 2018; Sabogal-Guáqueta et al., 2023). Building on these insights, the objective of this study was to comprehensively analyze the effects of nimodipine on activated microglia at the level of morphological and functional phenotypes, as well as their transcriptomic profile. We sought to identify Ca^2+^ dependent signaling cascades through which nimodipine may attenuate the pro-inflammatory shift in microglial function. The collected evidence suggests that nimodipine exerts distinct anti-inflammatory potential, which could be considered when expanding the field of clinical indication of nimodipine.

## Materials and Methods

### Brain slice preparations

Adult, C57BL/6 mice from both sexes (body weight: 18-24 g; n=6) were used in this study. Animals (Charles River Laboratories breed) were acquired from the Central Animal House of Biological Research Center, Szeged, Hungary. Standard rodent chow and tap water were supplied ad libitum. The animals were housed under constant temperature, humidity, and lighting conditions (23 °C, 12:12 h light/ dark cycle, lights on at 6 a.m.).

Coronal brain slices were prepared as previously reported (Frank et al., 2021). Briefly, C57BL/6 mice were deeply anesthetized with 5% isoflurane (in N2O:O2; 2:1), the animals were decapitated, 350 μm thick coronal brain slices were cut anterior to the bregma using a vibrating blade microtome (Leica VT1000S, Leica, Germany) and collected in ice-cold cutting aCSF (130 NaCl, 3. 5 KCl, 1 NaH2PO4, 24 NaHCO3, 1 CaCl2, 3 MgSO4, and 10 d-glucose in mM concentrations). After cutting, three to five slices were allowed to recover in carbogenated normal aCSF (130 NaCl, 3.5 KCl, 1 NaH2PO4, 24 NaHCO3, 3 CaCl2, 1.5 MgSO4, and 10 d-glucose in mM concentrations) for 30 min. Randomly selected slices were placed in an interface-type tissue chamber (Brain Slice Chamber BSC1, Scientifc Systems Design Inc., Ontario, Canada) and continuously perfused with carbogenated aCSF at a rate of 2.5 ml/min. Chamber temperature was maintained at 32°C using a dedicated proportional temperature control unit (PTC03, Scientifc Systems Design Inc., Ontario, Canada). Microglia in brain slice preparations were activated by mild oxygen-glucose deprivation (mOGD), achieved by superfusion of aCSF containing 5 mM D-glucose (reduced to 50%) for 45 min and transient hypoxia which was induced by the withdrawal of carbogen for 60 seconds three times at 15-minute intervals (Frank et al., 2024). Hypoxic episodes gave rise to SD (Frank et al., 2024). Brain slices were randomly assigned to a nimodipine-treated or a control group. The slices were bathed in nimodipine-enriched aCSF (10 μM final nimodipine concentration) for 30 min before and 45 min during mOGD, prior to histological processing.

### Maintenance and treatment of cell cultures

Primary cortical cell co-cultures were isolated from newborn rats, and microglial monocultures were derived from the co-cultures as previously described (Szabo and Gulya, 2013). Briefly, cerebrocortical tissue from newborn Sprague-Dawley rat pups (P1) was rapidly dissected, minced, and dissociated in 0.25% trypsin for 10 min at 37°C. The trypsin was then neutralized with Dulbecco’s modified Eagle’s medium (DMEM) containing 1 g/L D-glucose, 110 mg/mL Na-pyruvate, 4 mM L-glutamine, 3.7 g/L NaHCO3, 10,000 U/mL penicillin G, 10 mg/mL streptomycin sulfate, and 25 μg/mL amphotericin B and 15% heat-inactivated fetal bovine serum (FBS; Thermo Fisher Scientific, Carlsbad, CA, USA). After centrifugation at 1000g for 10 minutes at room temperature (RT), the pellet was resuspended, washed in 10 mL DMEM containing 10% FBS, and centrifuged again at 1000g and RT for 10 minutes. The final pellet was filtered through a sterile filter (100 μm pore size; Greiner Bio-One Hungary Kft., Mosonmagyaróvár, Hungary) to remove tissue fragments that had resisted dissociation. The cells were resuspended in 2 mL of the same solution and then seeded on poly-L-lysine-coated culture flasks (75 cm^2^; 10^7^ cells/flask) or poly-L-lysine-coated coverslips (15×15 mm; 2×10^5^ cells/coverslip) for immunocytochemistry or in poly-L-lysine-coated Petri dishes (60 mm × 15 mm; 10^6^ cells/Petri dish) for Western blot analysis and cultured at 37°C in a humidified air atmosphere supplemented with 5% CO2. The medium was changed the next day and every 3 days thereafter. After 7 days of culture, microglial cells in primary co-cultures (DIV7) were shaken on a platform shaker (120 rpm for 20 min) at 37°C, the supernatant was collected by centrifugation (3000g for 8 min at RT), resuspended in 2 mL DMEM/10% FBS, and seeded in the same medium either on poly-L-lysine-coated coverslips (15×15 mm; 2×10^5^ cells/coverslip) for immunocytochemistry or in poly-L-lysine-coated Petri dishes (60 mm × 15 mm; 10^6^ cells/dish) for Western blot analysis. The number of cells collected was determined in a Bürker chamber after trypan blue staining. DMEM/10% FBS was replaced the next day and then on the third and sixth day of subcloning (subDIV6).

Primary DIV6 co-cultures and microglia subDIV6 monocultures were challenged with lipopolysaccharide (LPS, dissolved in DMEM, 20 ng/mL in final concentration; Sigma, St. Louis, MO, USA) (Kata et al., 2016) and treated with nimodipine (dissolved in ethanol, 5 μM, 10 μM, 20 μM in final concentration; Sigma, St. Louis, MO, USA) on day 6. The following experimental conditions were established: (i) control (unchallenged, untreated), (ii) activated (challenged with LPS alone) for 24 h, (iii) treated with nimodipine alone (at 5, 10 and 20 μM final concentrations) for 24 h, and (iv) challenged with LPS at the presence of nimodipine (5, 10 and 20 μM final concentrations).

### Immunohistochemistry – cell cultures and brain slice preparations

For immunohistochemistry, we used a protocol described previously (Kata et al., 2017). Briefly, primary co-cultures (DIV7) and microglia (subDIV7) monocultures on poly-L-lysine-coated coverslips were used for immunohistochemistry. Cells were fixed in 4% formaldehyde in 0.05 M phosphate buffered saline (PBS, pH 7.4 at RT) for 5 min, then rinsed in 0.05 M PBS for 3 x 5 min. After permeabilization and blocking of nonspecific sites in 0.05 M PBS solution containing 5% normal goat serum (Sigma) and 0.3% Triton X-100 for 60 min at 37°C, the cells on the coverslips were incubated overnight at 4°C with the appropriate primary antibodies (Table S1) in 1% heat-inactivated bovine serum albumin (BSA; Sigma) and 0.3% Triton X-100. The cultured cells were washed 3 x 5 min at RT in 0.05 M PBS, then incubated with the appropriate Alexa Fluor fluorochrome-conjugated secondary antibody (Table S1) in the above solution, but without Triton X-100, for 2 h at RT in the dark. The cells on the coverslips were washed in 0.05 M PBS for 3 x 5 min at RT, and the nuclei were stained in 0.05 M PBS solution containing 1 mg/ml polyvinylpyrrolidone and 0.5 μl/ml 2-(4-amidinophenyl)-1H-indole-6-carboxamidine (DAPI; Thermo Fisher Scientific, Waltham, MA, USA). The coverslips were rinsed in distilled water for 5 minutes, air dried, and mounted on microscope slides in Vectashield mounting medium (Vector Laboratories, Burlingame, CA, USA).

Brain slices (350 µm) were rinsed with PBS, cryoprotected in 30% sucrose in PBS, embedded in paraffin, and sectioned (3 µm) on a rotation microtome (Leica RM 2235 Germany), as previously described (Frank et al., 2024). Sections were mounted on microscope slides, deparaffinized, and hydrated. Sections were then blocked with 10% normal goat serum (Sigma Aldrich, USA) for 1 hour at RT and incubated with rabbit anti-Iba1 primary antibody (Table S1) overnight at 4°C. Sections were rinsed with PBS, incubated with goat anti-rabbit secondary antibody (Table S1) at RT for 2 hours in the dark, then rinsed with PBS and distilled water. Sections were finally coverslipped with Fluoromount G (Thermo Fisher, USA., 00-4959-52) containing DAPI. Microscopic images of the parietal cortex and the striatum were taken with a LEICA DFC250 camera (Leica Microsystems Wetzlar GmbH, Wetzlar, Germany) attached to a fluorescence microscope (40× magnification, Leica DM LB2, Leica Microsystems CMS GmbH, Wetzlar, Germany).

### Western blot analysis – cell cultures

Cultured cells from primary co-cultures (DIV7) and microglial (subDIV7) monocultures were harvested with a rubber policeman, homogenized in 50 mM Tris-HCl (pH 7.5) containing 150 mM NaCl, 0.1% Nonidet P40, 0.1% cholic acid, 2 μg/ml leupeptin, 1 μg/ml pepstatin, 2 mM phenylmethylsulfonyl fluoride, and 2 mM EDTA, and centrifuged at 10,000g for 10 min at 4 ◦C. The pellet was discarded and the protein concentration of the supernatant was determined (Lowry et al., 1951). For quantitative analyses of microglial, neuronal, astrocyte, or oligodendrocyte immunoreactivity, 5-10 μg of protein was separated on a sodium dodecyl sulfate (SDS)-polyacrylamide gel (4-10% stacking gel/resolving gel), transferred to a Hybond-ECL nitrocellulose membrane (Amersham Biosciences, Little Chalfont, Buckinghamshire, England), blocked for 1 hour in 5% nonfat dry milk in Tris-buffered saline (TBS) containing 0.1% Tween-20 and incubated overnight with the appropriate primary antibodies (Table S1) as well as the internal control (mouse anti-GAPDH monoclonal antibody). After rinsing 5 times in 0.1% TBS-Tween-20, the membranes were incubated with the appropriate peroxidase-conjugated secondary antibodies (Table S1) for 1 hour and washed 5 times. The enhanced chemiluminescence method (ECL Plus Western blotting detection reagents; Amersham Biosciences) was used to detect immunoreactive bands according to the manufacturer’s protocol.

### In vitro phagocytosis assay – cell cultures

The fluid-phase phagocytic capacity of microglial cells in primary co-cultures (DIV7) and microglial (subDIV7) monocultures was determined by uptake of fluorescent microspheres (2 μm diameter; Sigma, St. Louis, MO, USA). Briefly, 1 μL per milliliter of a 2.5% aqueous suspension of fluorescent microspheres was added to the cultures, which were then incubated at 37°C for 60 minutes. The cells were then rinsed five times with 2 mL of PBS to remove any residual fluorescent microspheres bound to the dish or cell surface, and fixed with 4% formaldehyde in 0.05 M PBS (pH 7.4 at RT) (Szabo et al., 2013; Kata et al., 2017).

### Enzyme-linked immunosorbent assay (ELISA) – cell cultures

The concentrations of interleukin-10 (IL-10) (ER0033, FineTest) and tumor necrosis factor-α (TNF-α) (ER1393, FineTest) in both primary co-cultures (DIV7) and microglial (subDIV7) monoculture supernatants derived from the Western blot experiments were analyzed using an ELISA kit according to the manufacturer’s instructions. An automated microplate reader (Multiskan FC with the software SkanIt RE 5.0, Thermo Scientific) was used to measure optical density (OD) at 450 nm. The concentration of each sample was determined based on the optical density and the concentration of the standard. According to the manufacturer, the overall intra-assay and coefficient of variation were <8% for both IL-10 and TNF-a, and the inter-assay was <10% for both IL-10 and TNF-α.

### NEB mRNA-Library and next generation sequencing – cell cultures

Microglial monocultures were selected for gene expression analysis because the effect of nimodipine on microglial activation was essentially the same in both co-cultures and monocultures, and cell separation was not required for monocultures. Because 10 μM nimodipine was the lowest effective concentration of the drug on microglial activation, we focused the gene expression analysis on this treatment. Total RNA was isolated from the collected cells with RNeasy Mini Kit (74104 Qiagen) using 1 million cells/sample. Total RNA samples were quantified with Qubit 3.0 Fluorometer (ThermoFisher) and quality checked with Tape Station 4200 instrument using Agilent RNA ScreenTape (Agilent Technologies USA, Cat. No. 5067-5576).

NGS library preparation was carried out using the NEBNext Ultra™ II Directional RNA Library Prep Kit for Illumina (NEB #E7760) with NEBNext Poly(A) mRNA Magnetic Isolation Module (NEB #E7490). After mRNA enrichment, cDNA was generated where a second strand synthesis was done with the dUTP method to retain strand specificity. Finally, double-stranded cDNA was end-prepped and Illumina-specific adaptors were ligated to the cDNA fragments, followed by a final enrichment PCR.

Sequencing ready libraries were quality control checked by Tape Station 4200 instrument using D5000 ScreenTape (Agilent Technologies USA, Cat. No. 5067-5588). Next generation sequencing was carried out on NovaSeq X Plus sequencing system with NextSeq NovaSeq X Series 10B Reagent Kit (300 Cycle) chemistry (Illumina, Inc. USA, 20085594).

### Image analysis – cell cultures and brain slice preparations

Digital images were captured using a LEICA DFC250 camera (Leica Microsystems Wetzlar GmbH, Wetzlar, Germany) attached to a fluorescence microscope (Leica DM LB2, Leica Microsystems CMS GmbH, Wetzlar, Germany) and LAS X Application Suite (Leica Microsystems CMS GmbH, Wetzlar, Germany). To determine the purity of microglial cells, DAPI-labeled nuclei of Iba1-immunopositive cells were counted. In cell cultures, at least two coverslips from each experiment were analyzed, with approximately 20 randomly selected cells per coverslip in three separate experiments. In brain slices, five cells from both cortex and striatum were analyzed from six coverslips per group. Microglial cell silhouettes were obtained by converting the raw digital files of Iba1-immunoreactive cells captured by fluorescence microscopy into binary files using Adobe Photoshop CS3 software (Adobe Systems, Inc., San Jose, CA, USA). After binary conversion, cell perimeter and area of individual cells were measured in ImageJ (National Institutes of Health, Bethesda, MD, USA) and the TI reflecting the degree of process extension was calculated as follows: perimeter2/area* 4π as previously described (Fujita et al., 1996). A total of 420 cell silhouettes were analyzed in this study.

To measure phagocytic activity, cells were labeled with phagocytosed microbeads. The microbeads in the cytoplasm were counted. Twenty non-overlapping random fields were captured using a fluorescence microscope with a 40x objective (Leica DMLB epifluorescence microscope), and the microbead load of a total of 100 cells in each culture was evaluated using the ImageJ cell counter plugin (National Institutes of Health, Bethesda, MD, USA). The mean microbead count of 100 cells was taken as a single value representing each culture.

Gray-scale digital images of the immunoblots were obtained by scanning the autoradiographic films with a desktop scanner (Epson 430 Perfection V750 PRO; Seiko Epson Corp., Suwa, Japan). Images were scanned and processed with identical settings to allow comparison of blots from different samples. The bands were outlined and analyzed by densitometry using ImageJ (National Institutes of Health, Bethesda, MD, USA). Immunoreactive densities of equally loaded lanes were quantified as we previously reported (Kata et al., 2016, 2017). Samples were normalized to the densities of internal controls (GAPDH) and presented as % of controls.

### Statistical analysis

Statistical comparisons for microglial morphology, Western blot, phagocytic activity, and ELISA assay were performed using GraphPad Prism 8.0 (Windows, GraphPad Software, San Diego, California USA, www.graphpad.com). Normality of data distribution was determined by the Shapiro-Wilk test. In case of normal distribution of data, one-way analysis of variance (ANOVA) followed by Tukey’s multiple comparison test was used for statistical analysis, and data were presented as mean±SD in bar graphs. In case of non-normal distribution of data, Kruskal-Wallis test followed by Dunn’s multiple comparison was used and data were presented in box plots. The significance level was set at p < 0.05.

For the processing of RNA-Sequencing data, the quality of the pair-ended reads was performed using FastQC (0.12.1) (Andrews, 2010), while adapters and low-quality bases were trimmed using FASTP (0.23.4) (Chen et al., 2018). The filtered reads were aligned to the Rattus norvegicus genome (NCBI: GRCr8) using the splice-aware aligner, STAR (2.5.2b) (Dobin et al., 2013), and quantified with FeatureCounts (v.2.0.6) (Liao et al., 2014). For the downstream analysis, only genes with a CPM (counts per million) greater than 2 in at least three samples were retained. The filtered samples were then normalized using the TMM (Trimmed Mean of M-values) method implemented in edgeR’s calcNormFactors (v.4.0.16) function (Robinson et al., 2010). Differential gene expression analysis was performed using limma-voom (Law et al., 2014).

PCA and hierarchical clustering were used to examine the similarities between samples and identify potential outliers. Differently expressed genes (DEGs) were identified for the following comparisons: (i) LPS vs. Control, (ii) LPS+nimodipine vs. Control, and (iii) LPS+nimodipine vs. LPS. In each comparison, DEGs were defined as those with a log2 fold change greater than 1 and an adjusted p-value < 0.05.

To assess the effects of different treatments on gene expression, we selected genes from various pathways and analyzed their expression changes. Log Fold Change and p-values were used to evaluate the differential expression of these genes across comparisons. We identified significantly upregulated and downregulated genes based on three significant levels for these pathways. Additionally, heatmaps were generated for each pathway to provide a detailed view of expression patterns. The complete results of this analysis, including all significant genes, are available in the supplementary materials.

## Results

### Nimodipine alters the morphological phenotype of activated microglia

First, we recapitulated and extended our findings made previously in metabolically challenged live brain slice preparations (Frank et al., 2024). The morphological phenotype of microglia corresponds to the state of microglial activation, with a ramified morphology associated with the quiescent state and an ameboid shape indicating activation. Microglia acquired an apparent amoeboid, activated morphological phenotype upon mOGD challenge (TI 2-4) (Fig. 1). Nimodipine (at a concentration of 10 μM) partially restored the ramified morphology in both the cerebral cortex and striatum in the mOGD-challenged brain slices (TI 6-8) (Fig. 1).

**Figure 1.**
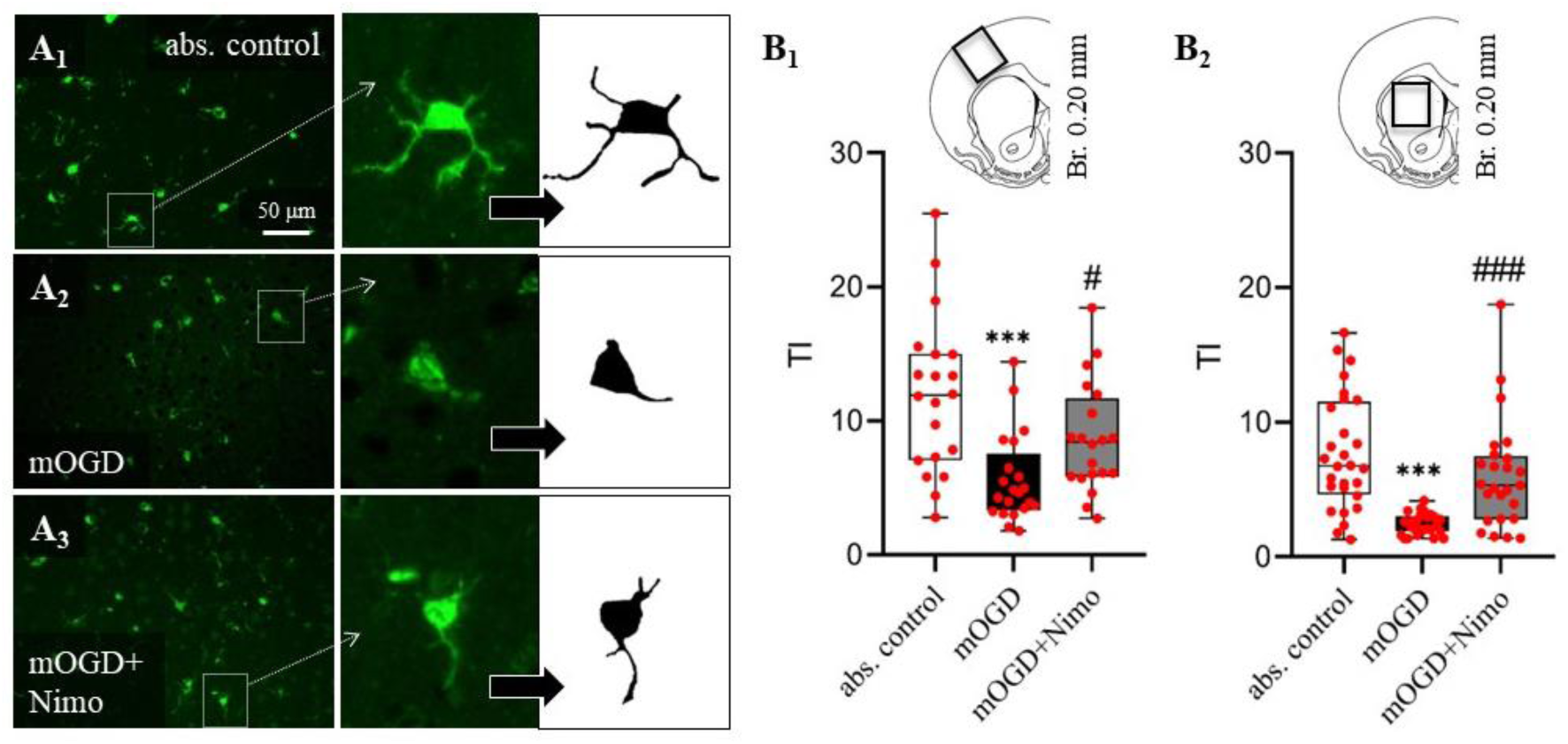
Quantitative analysis to evaluate the effect of nimodipine on microglial morphology in live brain slice preparations. **A,** Fluorescence immunohistochemistry images show Iba1-labeled microglia in absolute control brain slices (A_1_), in slices exposed to mild oxygen glucose deprivation (mOGD) (A_2_), and to mOGD in combination with nimodipine (mOGD+Nimo) (A_3_) in the cerebral cortex. Binary silhouettes of individual Iba1-positive microglia show the arborization of the cells. **B,** Transformation index (TI) calculated to quantify the degree of arborization (decreasing TI corresponds to an amoeboid shape and thus increasing activity) in the cortex (B_1_) and striatum (B_2_). Data are presented as min to max in box plots, with individual cells given by spheres (n=30). Normality of the data distribution was determined by the Shapiro-Wilk test (B_1_: p=0.022; B_2_: p=<0.001). Statistical analysis was based on Kruskal-Wallis test (B_1_, p<0.001***; B_2_, p<0.001***) followed by Dunn’s post hoc test (p<0.001*** vs. abs. control; p<0.05^#^ and p<0.001^###^ vs. mOGD).

To explore nimodipine’s direct effect on microglia, microglial cell cultures were used. In the untreated co-cultures, microglia typically exhibited a quiescent morphology, with small pseudopodia (Fig 2A-B). In the microglial monocultures, the cells were slightly more ramified with TI ∼4 (Fig. 2C-D). Activation with LPS significantly decreased TI in both co-cultures and monocultures, corresponding to an amoeboid morphological phenotype (Fig. 2). When applied to non-activated cultures, nimodipine (5, 10, 20 μM) induced the formation of numerous microspikes (filopodia) in both co-cultures and monocultures (Fig. 2A, C), but did not result in significantly increased TI values (Fig. 2B, D). The addition of nimodipine to LPS-activated cultures increased microglial TI close to control levels, especially at 10-20 µM concentrations, and resulted in a more quiescent phenotype in both co-cultures and monocultures (Fig. 2). This indicates that nimodipine at concentrations of 10 and 20 µM effectively counteracted the morphological changes characteristic of LPS-induced microglial activation.

**Figure 2.**
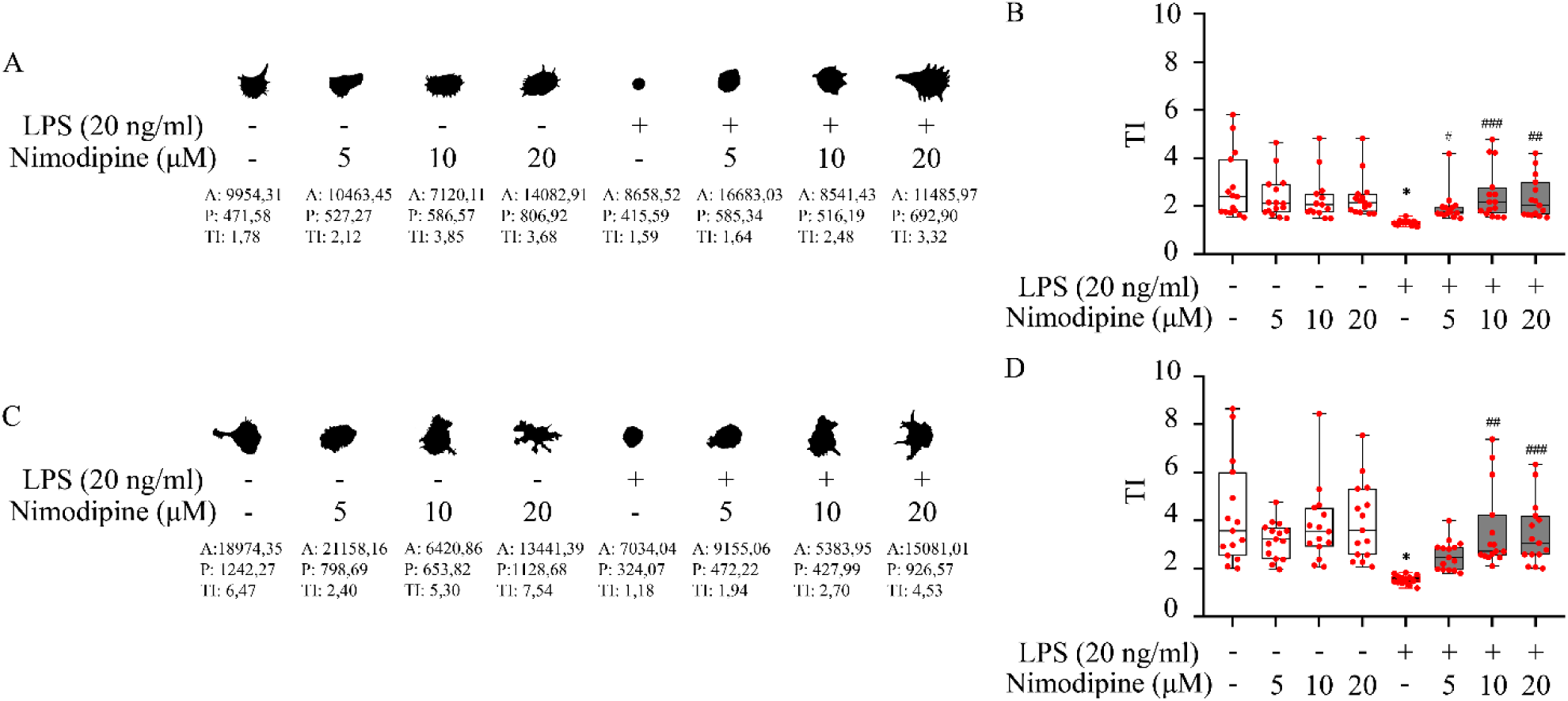
Quantitative analysis to evaluate the effect of nimodipine on microglial morphology in co-cultures (A-B) and microglia monocultures (C-D). **A,** Iba1-positive cell masks representative of microglia in co-cultures (DIV7). **B,** Microglial transformation index (TI) calculated for each experimental condition in co-cultures. **C,** Iba1-positive cell masks representative of microglia in mono-cultures (subDIV7). **D,** Transformation index (TI) calculated for each experimental condition in microglia mono-cultures. In A and C, the values are given below the masks: A, area; P, perimeter; TI, transformation index. In B and D, data are presented as min to max in box plots; red spheres represent individual values in each condition (n=15). Normality of the data distribution was determined by the Shapiro-Wilk test (B: p<0.001; D: p<0.001). Data were analyzed by Kruskal-Wallis test (B, p<0.001***; D, p<0.001***) followed by Dunn’s multiple comparison (p<0.05* vs. absolute control; p<0.05^#^, p<0.01^##^, p<0.001^###^ vs. LPS alone).

### Nimodipine reduces microglial Iba1 levels after activation

Microglial Iba-1 expression was taken as an indicator of activation (Ito et al., 1998). Western blot analysis showed that Iba-1 levels were increased in LPS-treated co-cultures (169.0±33.7 % of control) and monocultures (213.4±29.7 % of control) compared to the non-activated control condition (Fig. 3). Nimodipine did not alter Iba1 levels in non-activated cultures. In contrast, in LPS-activated co-cultures, the addition of nimodipine resulted in a concentration-dependent reduction of Iba1 expression that reached significance at 10 and 20 µM nimodipine concentrations (103.7±8.3 and 64.9±15.9 vs. 169.0±33.7 % of control, LPS + 10 and 20 µM nimodipine vs. LPS) (Fig. 3B). In microglia monocultures, nimodipine reduced Iba1 levels to control at all three concentrations used (95.3±18.1, 108.3±20.1 and 77.2±8.5 vs. 213.4±29.7 % of control, LPS + 5, 10 and 20 µM nimodipine vs. LPS) (Fig. 3D).

**Figure 3.**
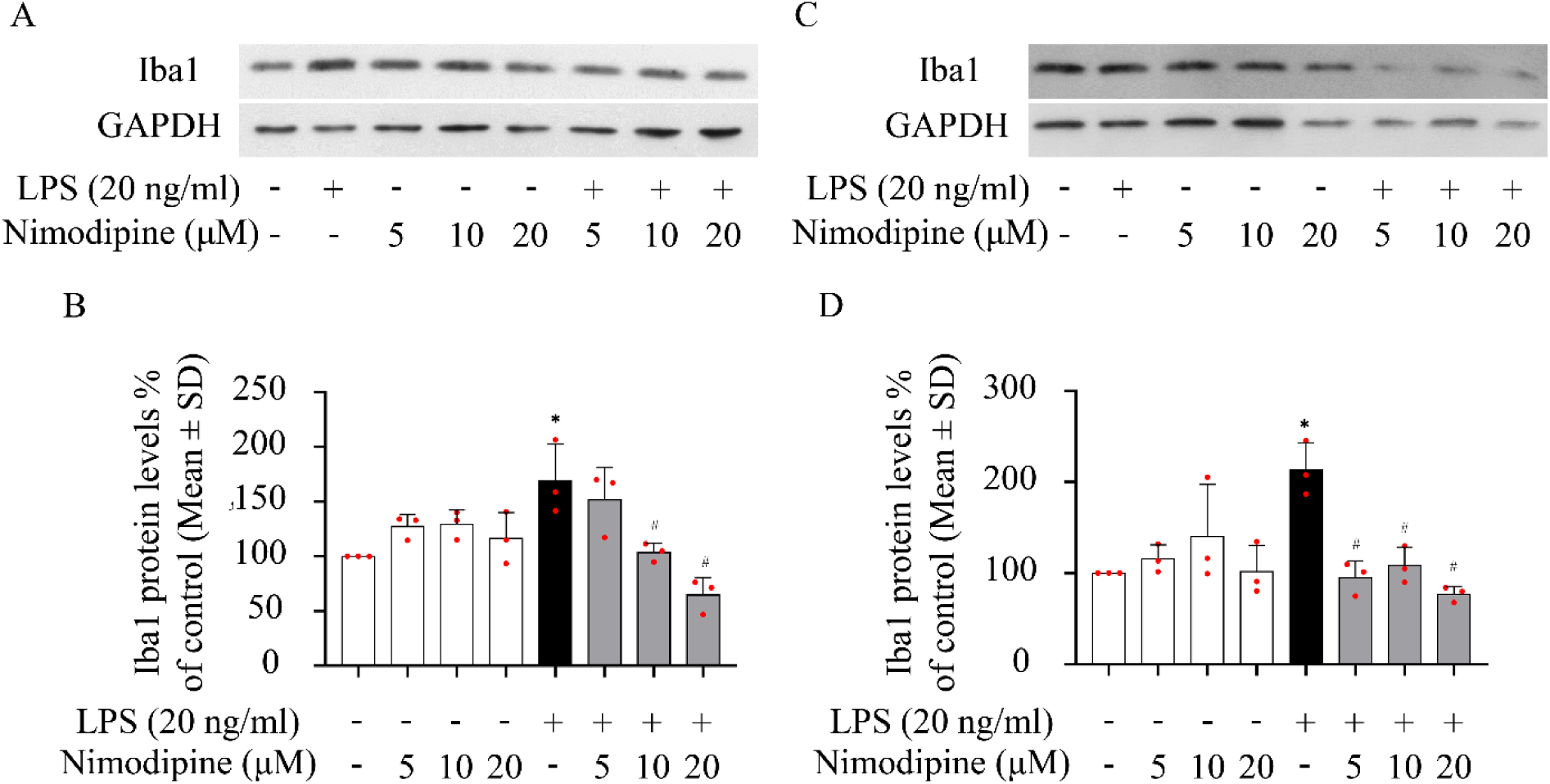
Quantitative analysis to detect the effect of nimodipine on microglial Iba1 protein levels in co-cultures (A-B) and microglial monocultures (C-D). **A,** Representative Western blot images for Iba1 and GAPDH, used as an internal control, obtained from co-cultures. **B,** Iba1 protein levels quantified in co-cultures. **C,** Representative Western blot images for Iba1 and GAPDH from microglial monocultures. **D,** Iba1 protein levels quantified in microglia monocultures. In A and C, images were scanned and processed with identical settings to allow comparisons between Western blots from different samples. In B and D, data are integrated optical density values relative to control and are presented as mean±SD (n=3). Red spheres indicate individual values in each group. Normality of data distribution was determined by Shapiro-Wilk test (B: p=0.970; D: p=0.284). Data were analyzed by one-way analysis of variance (ANOVA) (B, f=7.802, p<0.001***; D, f=7.123, p<0.001***) followed by Tukey’s multiple comparison (p<0.05* vs. absolute control; p<0.05^#^ vs. LPS alone).

### Nimodipine reduces phagocytic activity of activated microglia

Non-activated microglia exhibited low levels of phagocytosis, engulfing only 0.9±0.3 microbeads per cell in co-cultures and 4.7±1.0 in monocultures. Nimodipine applied to non-activated cultures had no significant effect on phagocytosis, with the number of phagocytosed microbeads remaining low (co-cultures: 1.19±0.2; monocultures: 4.21±1.2 microbeads per cell). As expected, the LPS challenge significantly increased microglial phagocytic activity. On average, LPS-activated microglia accumulated 3.5±0.3 microbeads per cell in co-cultures and 11.5±1.2 microbeads per cell in monocultures. The addition of nimodipine significantly inhibited phagocytosis, which returned to near control levels in both co-cultures (1.29±0.15 microbeads per cell) and in monocultures (4.63±1.14 microbeads per cell) (Fig. 4).

**Figure 4.**
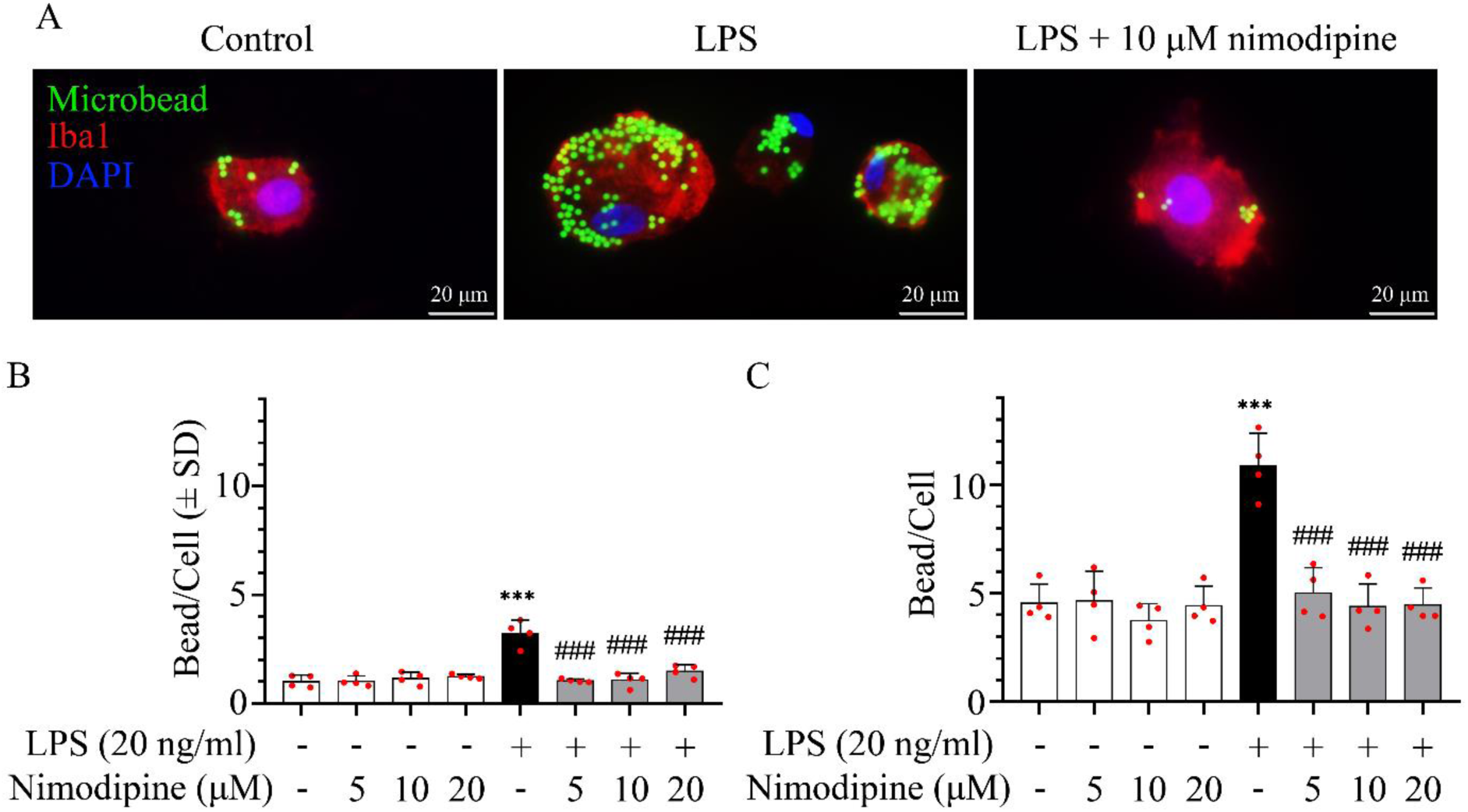
Evaluation of the effect of nimodipine on microglial phagocytic activity as visualized by fluorescent microbeads co-localized with Iba1 protein. **A,** Representative fluorescence microscopic images of Iba1-labeled microglia in monocultures. Fluorescent microbeads in their cytoplasm indicate increased phagocytic activity after LPS activation, which was counteracted by nimodipine. **B,** Phagocytotic activity of microglia quantified in co-cultures. **C,** Phagocytotic activity quantified in microglial monocultures. In B and C, data are presented as mean±SD; red spheres represent individual cultures (mean of 100 cells analyzed in each culture) in each group. Normality of data distribution was determined by Shapiro-Wilk test (B, p= 0.259; C, p=0.216). Data were analyzed by one-way analysis of variance (ANOVA) (f=49.640, p<0.001***) followed by Tukey’s multiple comparison (p<0.001*** vs. absolute control; p<0.001^###^ vs. LPS alone).

### Nimodipine reduces the secretion of TNF-α by activated microglia

Activated microglia express both pro- and anti-inflammatory cytokines. Here, we tested whether nimodipine regulates cytokine release, and evaluated the levels of the pro-inflammatory cytokine TNF-α and the anti-inflammatory cytokine IL-10 in the culture media. The basal level of IL-10 was 146.98±24.19 pg/ml in co-culture media and 144.5±7.89 pg/ml in monoculture media. LPS at the low concentration used (20 ng/ml) did not significantly alter IL-10 protein expression (180.26±0.87 and 123.25±65.59 pg/ml, co-cultures and monocultures, respectively). Nimodipine had no effect on IL-10 levels in either non-activated or activated cultures.

The level of TNF-α was 5.37±1.64 pg/ml in non-activated co-cultures and 6.74±3.43 pg/ml in monocultures (Fig. S1). Low dose LPS activation did not significantly change the TNF-α concentration measured at 24 h (peak values had been previously measured at 6 h, Kata et al., 2016), despite a visible tendency to increase both in co-cultures (from 5.37±1.64 to 15.52±4.95 pg/ml) and in monocultures (from 6.74±3.43 to 33.35±17.26 pg/ml). Again, nimodipine had no effect on non-activated cultures. However, nimodipine at a concentration of 20 μM reduced TNF-α concentration in LPS-activated co-cultures to the control levels (1.61±0.41 pg/ml). In LPS-activated monocultures, nimodipine produced a stepwise reduction in TNF-α levels along its increasing concentration (27.76±19.54, 23.25±12.27 and 13.61±10.26 pg/ml, 5, 10 and 20 µM), although without statistical significance.

### LPS markedly alters gene expression indicating pro-inflammatory microglial phenotype

Next-generation RNA sequencing was conducted on microglial monocultures. Principal component analysis (PCA) of the transcriptome data showed that replicates of the control group clustered closely together, and the LPS- and LPS+nimodipine-treated samples were clearly separated from the control. The LPS and LPS+nimodipine samples were heterogeneous and did not form distinct clusters, revealing a predictably weaker effect of nimodipine on gene expression than that of LPS (Fig. S2). The PCA results were reflected in the unsupervised hierarchical clustering of the samples (Fig. S2). In total, 13,866 genes were identified by next-generation RNA sequencing in primary microglial monocultures, of which 9,720 were differentially expressed (DEGs) by LPS challenge alone or in combination with nimodipine treatment (Fig. 5).

**Figure 5.**
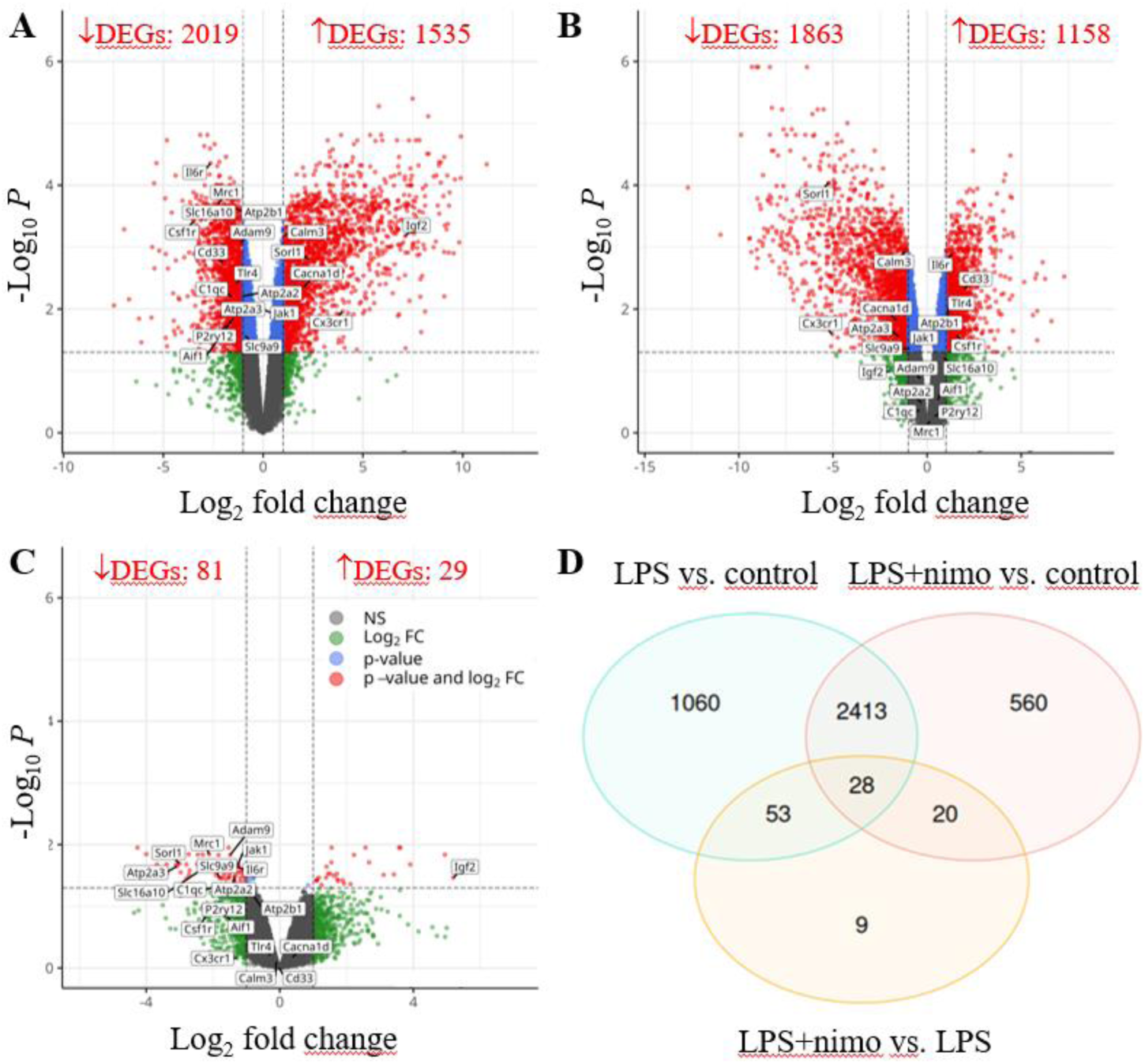
Activation with lipopolysaccharide (LPS) and treatment with nimodipine (nimo) cause alterations in rat primary microglia transcriptome. Volcano plots depict Log_2_ fold changes and -Log_10_ of the adjusted p-value per gene comparing LPS vs. control (**A**), LPS+nimodipine treatment vs. control (**B**) and LPS+nimodipine treatment vs. LPS alone (**C**). Venn diagram shows the overlap of differentially expressed genes (DEGs) in the three experimental conditions (**D**).

Next, we analyzed gene expression patterns linked to distinct functions of activated microglia. LPS is specifically recognized by toll-like receptor 4 (TLR4) receptors located on the microglial plasma membrane (Lenhardt et al., 2003). Indeed, the expression of the TLR4 gene (Tlr4) was significantly upregulated in the LPS-challenged microglia. Downstream of TLR4, the expression of several genes encoding adaptor proteins of TLR4 signaling cascades (Zhou et al., 2020) were all upregulated (e.g. Myd88, Ticam1, Ticam2) (Fig. S3).

Further, LPS considerably enhanced cytokine signaling in the primary microglial cultures. Specific genes encoding interleukins and chemokines were upregulated (e.g. Il7, Il11, Il16, Il18, Ccl20, Cxcl6, Cxcl16), while others were downregulated (e.g. Il17d, Il34, Ccl19, Cxcl12, Cxcl14) (Fig. S4). The gene expression of a group of cytokine receptors was also significantly upregulated (e.g. Il1r1, Il6r, Il7r, Il23r, Tnfrsf1b, Tnfrsf8, Tnfrsf25, Tnfrsf26) (Fig. S4). Some of the markers previously recognized to represent M1 polarization (Jurga et al., 2020) were expressed at significantly higher levels (e.g. Ccl20, Cd86, Il18), whereas gene expression typical for M2 polarization (e.g. Il10, Ccl2, Ccl22, Cxcl13, Arg1) (Jurga et al., 2020) was not altered in the LPS-activated cultures, suggesting a predominantly pro-inflammatory microglial phenotype (Fig. S4).

In addition to the widely appreciated cytokine response of microglia to pathogen- or damage-associated molecular patterns, microglia release complement components and express complement receptors, which appear to contribute to pathological mechanisms in neurodegenerative disorders (Chen et al., 2022). Of the first elements of the classical complement cascade, the LPS challenge upregulated genes of the complement component C1q (C1qa and C1qc) and downregulated complement components C1r and C1s. Downstream the classical pathway, complement components C4a and C4b were also expressed at significantly lower levels. On the other hand, the expression of complement receptor genes C3ar1 and C5ar1 were markedly upregulated by LPS (Fig. S5).

Consistent with the increased phagocytic activity in response to LPS shown here (Fig. 4), the expression levels of phagocytic receptor genes (Podleśny-Drabiniok et al., 2020; Thomas et al., 2022) were significantly increased (e.g. Mertk, Trem1, Cd14, Cd33, Cd68, Ms4a4a), except for Sorl1, which was downregulated (Fig. S6). Autophagy, a catabolic mechanism to degrade dysfunctional proteins and organelles may suppress inflammation (Jin et al., 2018). The expression of genes associated with macroautophagy (e.g. Mtor, Ulk1, Atg6l, Map1lc3b or Sqstm1), however, were not affected by LPS. Yet, the genes of some proteins that regulate autophagy (e.g. Wdfy3 and Dnajc16) were expressed at a higher level due to LPS.

Finally, as recently reported (Sabogal-Guáqueta et al., 2023), LPS induced a metabolic shift in cultured microglia. In general, genes encoding glycolytic enzymes were upregulated, whereas genes of some enzymes of the TCA cycle were downregulated (Fig. S7). In particular, hexokinase genes (Hk2 and Hk3) were expressed at higher levels, in line with previous reports (Sabogal-Guáqueta et al., 2023), indicating the potential for increased glucose phosphorylation.

### LPS modifies intracellular calcium homeostasis with distinct action of nimodipine

Because microglia respond to stimulation with intracellular Ca^2+^ accumulation, and nimodipine is a calcium channel blocker, we analyzed how proteins implicated in intracellular Ca^2+^ homeostasis are altered by LPS and nimodipine.

Calcium influx across the plasma membrane in microglia is mediated by VGCCs, ligand-gated cation channels, transient receptor potential (TRP) cation channels, and Ca^2+^ release-activated Ca^2+^ channels (Fig. S8). The gene expression of many of these pathways of Ca^2+^ movement was altered by LPS. In particular, some of the genes of the pore-forming α subunit of VGCCs were remarkably downregulated (Cacna1d, Cacna1e, Cacna1g, Cacna1h). Among the ligand-gated cation channels, the expression of the genes encoding the 3A subunit of the NMDA receptor (Grin3a) and subunits 1 and 2 of the AMPA receptor (Gria1 and Gria2) was decreased, while the expression of the gene encoding the P2RX4 purinergic receptor (P2rx4) was increased. Regarding TRP channels, the expression of the gene of TRPM2 (Trpm2) was upregulated. Finally, the gene Orai1, which encodes a Ca^2+^ release-activated Ca^2+^ channel, was not affected. Nimodipine counteracted the LPS effect only on the expression of the Gria1 gene, the expression of other genes of plasmalemmal Ca^2+^ channels were not affected (Table 1).

**Table 1.**
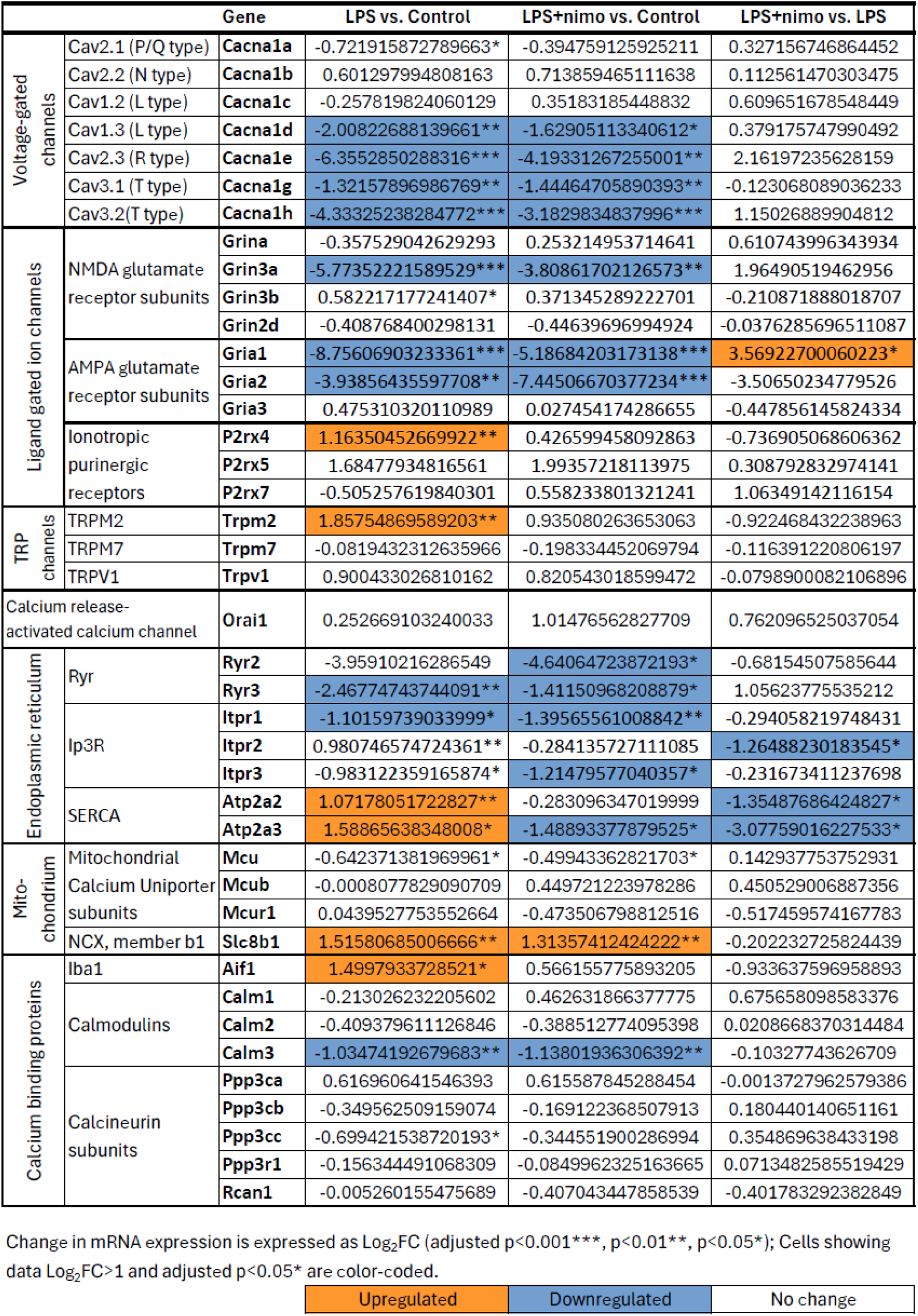
Changes in gene expression of calcium channels and calcium binding proteins.

The endoplasmic reticulum (ER) is the major Ca^2+^ store in cells and is equipped with ligand-gated ryanoid receptors (RyRs) and inositol triphosphate receptors (IP3Rs) to mediate Ca^2+^ release upon activation (Fig. S8). In addition, Ca^2+^ pumps (SERCA) ensure Ca^2+^ uptake into the ER against the Ca^2+^ concentration gradient at the expense of ATP. The expression of all these genes was remarkably modulated by LPS treatment. Comparison of LPS-challenged microglia with the control showed that RyR genes (Ryr2-3) along with the IP3R genes (Itpr1-3) were downregulated. In contrast, the expression of the IP3R gene Itpr2 was upregulated by LPS. Genes encoding SERCA (Atp2a2 and Atp2a3) were remarkably upregulated by exposure to LPS. Interestingly, the effect of nimodipine treatment was apparent only on the genes upregulated by LPS. Comparing the LPS+nimodipine group with the LPS alone group, Itpr2, Atp2a2, and Atp2a3 were downregulated by nimodipine treatment (Table 1).

Among the Ca^2+^ binding proteins, the gene of Iba1 (Aif1) was upregulated by LPS (Table 1), consistent with protein levels measured with Western blot analysis here (Fig. 3). Although nimodipine showed a tendency to counteract the LPS effect on Aif1 expression, the effect was not significant at mRNA level (Table 1) in contrast with a significant reduction at protein level (Fig. 3). The expression of the Calm3 gene coding for the calcium binding protein calmodulin 3 was markedly decreased by LPS, but nimodipine did not change the gene expression of calmodulins (Table 1).

### Nimodipine exerts complex action on microglial immune response, acting at multiple sites

In our primary microglial monocultures, nimodipine administered with LPS altered the expression of 110 genes compared to LPS alone (Figs. 5-6). Of these DEGs, 29 were upregulated and 81 were downregulated. Remarkably, nimodipine had opposite effects to LPS (Fig. 6). Screening of these 110 genes revealed that at least 20 were associated with microglial immune response, 7 with cell adhesion, and 2 with autophagy regulation (Fig. 6), in addition to 4 involved in intracellular Ca^2+^ homeostasis shown above. Downregulated DEGs involved in immune function included genes of toll-like receptor 9 (Tlr9) that was upregulated by LPS alone, in agreement with previous reports (Olson and Miller, 2004), and a negative feedback regulator of TLR signaling (Zc3h12d). The expression level of cytokine receptors interleukin-6 receptor (Il6r) and a TNF receptor (Tnfrsf8) increased by LPS were also downregulated due to nimodipine treatment, along with the immune suppressor modulator fibrinogen-like protein 2 (Fgl2). Concerning the complement system, complement C1q (C1qc) and a complement receptor of complement fragment C3b (Vsig4) were downregulated by nimodipine opposing LPS effect. Finally, the expression of the MHC class II transactivator CIITA (Ciita) and CD36 (Cd36) also decreased at the presence of nimodipine. Upregulated DEGs associated with immune function were somatostatin 4 receptor (Sstr4), proposed to regulate microglial activation and phagocytosis (Sandoval et al., 2019; Silwal et al., 2022), and an a-disintegrin and metalloproteinase with thrombospondin motifs (Adamts12) (Table 2; Fig. 6).

**Figure 6.**
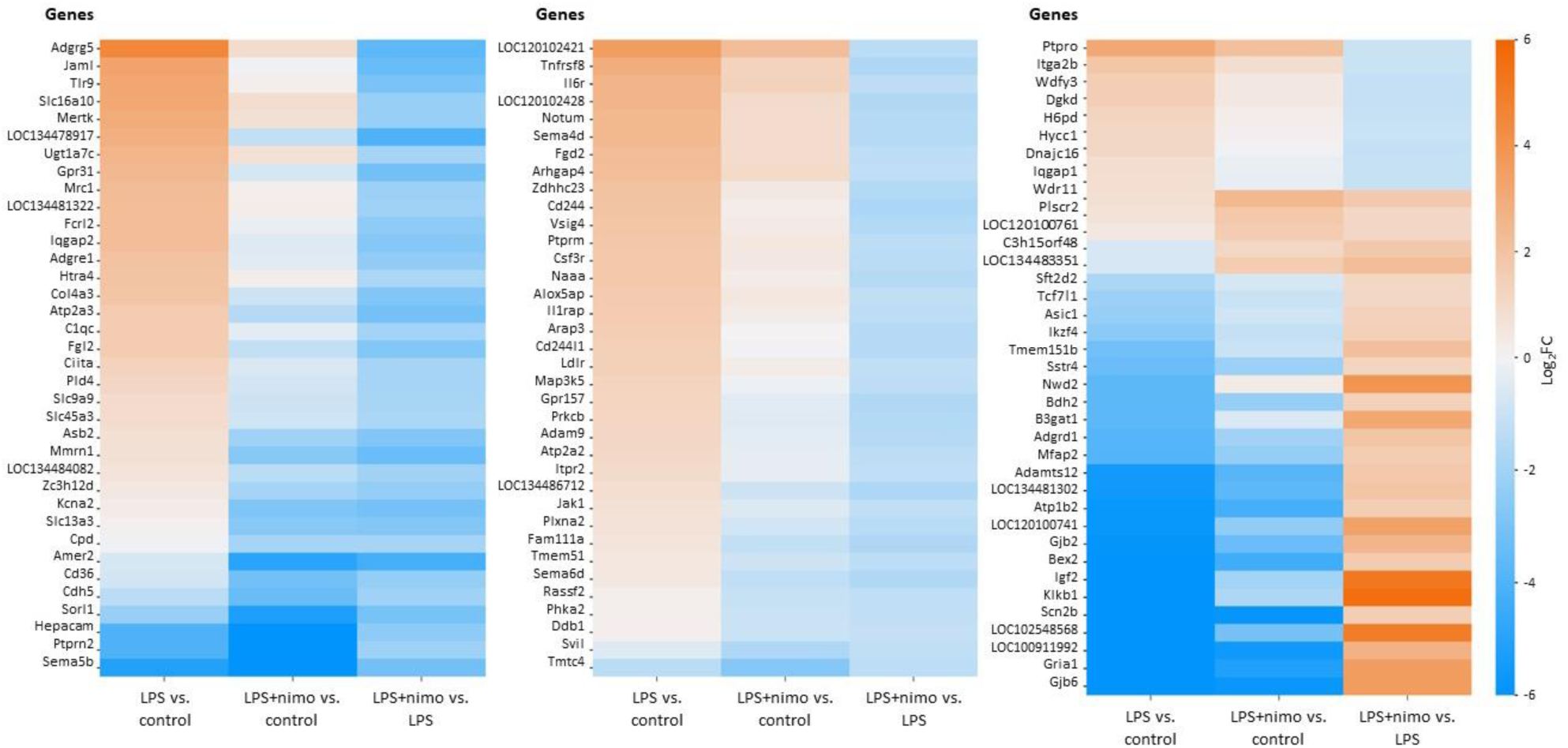
Heat maps of differentially expressed genes (DEGs) selected for significant change by nimodipine treatment (LPS+nimodipine) compared to LPS challenge alone (selection criteria: -1>Log_2_FC>1 and adj.p<0.05)

**Table 2.**
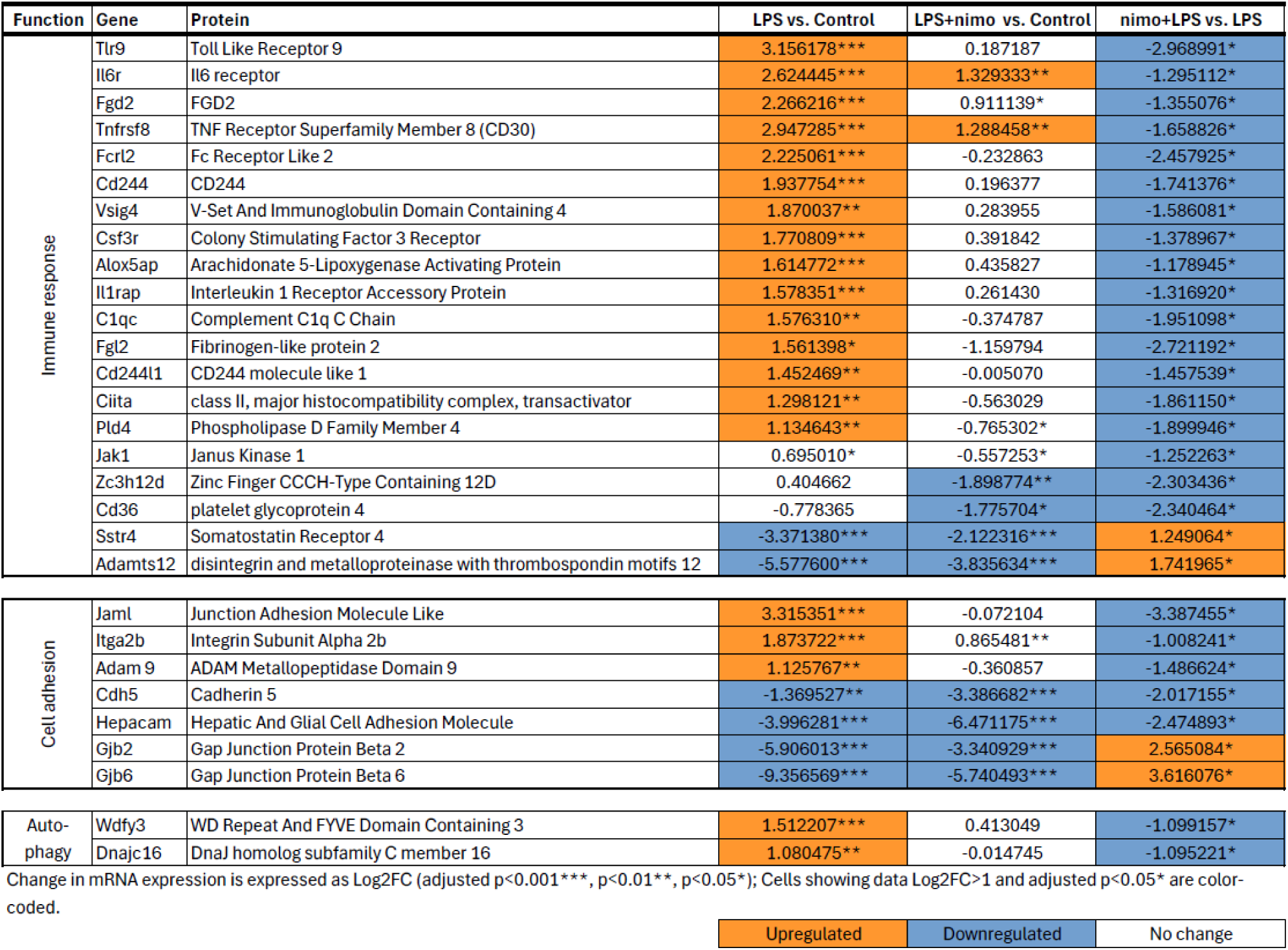
Selected differentially expressed genes (DEGs) by nimodipine treatment, grouped by function.

Among the DEGs associated with cell adhesion, a member of the junctional adhesion molecule transmembrane protein family (Jaml), an a-disintegrin and metalloproteinase (Adam9), and an integrin cell adhesion receptor (Itga2b) were downregulated in nimodipine-treated cultures, counteracting LPS effect. In contrast, genes of gap junction proteins beta 2 and beta 6 (Gjb2 and Gjb6, respectively) were upregulated, opposing LPS. Finally, genes of the autophagy regulators Wdfy3, which bridges cargo to the molecular machinery that builds autophagosomes (Filimonenko et al., 2010; Napoli et al., 2021), and Dnajc16, which determines the size of autophagosomes (Yamamoto et al., 2020), were downregulated by nimodipine, counteracting LPS effect (Table 2; Fig. 6).

Finally, functional enrichment analysis confirmed the positive effect of nimodipine on intercellular connections (e.g., connexin complex, gap junction, intercellular transport), and elucidated the negative effect on the activity of several enzymes implicated in intracellular signal transduction (e.g., kinases and transferases), in addition to Ca^2+^ trafficking of the ER (Fig. 7).

**Figure 7.**
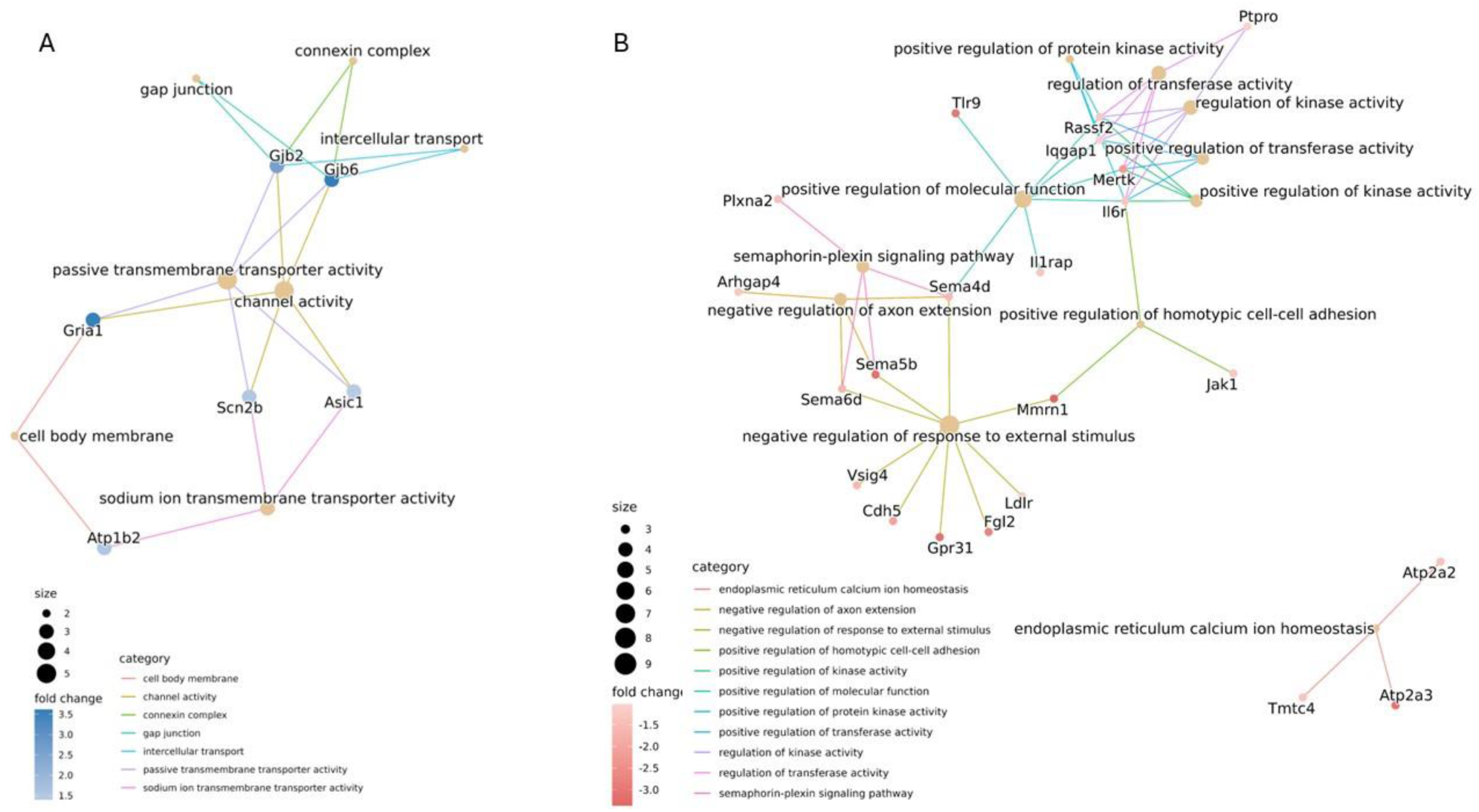
Functional enrichment analysis results are displayed in gene-concept network plots (cnet plots) to illustrate the effect of nimodipine on biological processes (BP). Upregulated (A) and downregulated (B) functions are shown by linkages of genes and gene ontology (GO) terms (LPS+nimodipine vs. LPS alone).

## Discussion

In this study, exposing live brain slice preparations to mOGD or challenging microglial cultures with LPS resulted in microglial activation, which was clearly indicated by a shift toward ameboid morphology (Figs. 1-2), and, in cell cultures, by increased phagocytic activity (Fig. 4). The shift in microglial phenotype was underscored by complex changes in gene expression (Fig. 5). Importantly, the addition of nimodipine to the activated brain slice preparations and microglial cultures suppressed microglial activation (Figs. 1-4), caused change in the expression of a distinct group of genes associated with immune response (Figs. 5-6, Tables 2-3), and thus delivered two main messages. First, Ca^2+^ influx through LVGCCs plays an important role in microglial activation, by modulating gene expression. Second, the LVGCC blocker nimodipine confers partial protection against microglial activation by modulating distinct elements of the immune response, and may offer benefit as an anti-neuroinflammatory agent.

There is a consensus that microglial activation is associated with increased intracellular Ca^2+^ concentration in response to a variety of danger signals (Hopp et al., 2020), including the widely used activator LPS (Hoffmann et al., 2003). Less is known about which pathways of Ca^2+^ movement are involved and how an inflammatory stimulus redirects transmembrane microglial Ca^2+^ currents. Here we show that the expression of the purinergic P2X4 receptors and TRPM2 channels was upregulated by LPS, suggesting that these two pathways may be the predominant Ca^2+^ influx pathways sustained during microglial activation. Consistent with our data, the microglial P2X4 receptors activated by elevated extracellular ATP concentrations open to allow Ca^2+^ influx and initiate an inflammatory response (Suurväli et al., 2017), and are upregulated in various brain and spinal cord pathologies involving neuroinflammation (Montilla et al., 2020). Similarly, LPS/IFNγ stimulation of microglia has been shown to induce TRPM2-mediated Ca^2+^ signaling (Miyake et al., 2014), and TRPM2 channels have been identified as part of a feedback loop that promotes pro-inflammatory polarization (Raghunatha et al., 2020). In contrast, we found that the expression of VGCCs and glutamate-gated cation channels (NMDA and AMPA receptors) was downregulated in response to LPS, implying reduced Ca^2+^ conductance through these channels, which may be compensative. Our data are consistent with those previously reported showing that cultured microglia express the Cav1.2 and Cav1.3 proteins of LVGGCs (Espinosa-Parrilla et al., 2015). However, we did not observe an upregulation of these genes, in contrast to the increased Cav1.2 protein levels quantified earlier by Western blotting, after LPS/TRP stimulation of BV2 microglia (Espinosa-Parrilla et al., 2015). The discrepancy in the results may be due to the different model systems (primary microglial cultures used here versus an immortalized cell line used previously) or post-translational modification of Cav proteins (Szymanowicz et al., 2024) to cause difference in the levels of mRNA evaluated here versus protein concentration assessed previously (Espinosa-Parrilla et al., 2015). Importantly, although Cacna1c and Cacna1d gene expression was reduced by LPS in our preparations (Table 1), our results imply that the LVGCC proteins Cav1.2 and Cav1.3 (encoded by Cacna1c and Cacna1d, respectively) must have been abundant enough in the plasmalemma to mediate Ca^2+^ influx relevant to microglial activation and to serve as the target of nimodipine to produce the observed effects.

In addition to the lower expression of the above mentioned plasmalemmal Ca^2+^ channels, Ca^2+^ release from the ER must also have been inhibited during activation, as evidenced by the downregulation of several ryanoid and IP3 receptor genes. At the same time, active Ca^2+^ transport into the ER appeared to be augmented by the upregulation of SERCA genes (Atp2a2 and Atp2a3), in agreement with recent findings of others focusing on SERCA2b (Morales-Ropero et al., 2021). These changes suggest that Ca^2+^ storage in the ER is favored during microglial activation, presumably to counterbalance the intracellular Ca^2+^ load through membrane channels in response to the inflammatory stimulus. This concept seems to be supported by the observation that the SERCA genes were downregulated when nimodipine, a Ca^2+^ channel blocker was added to our preparations (Table 1). Nimodipine must have reduced the overall Ca^2+^ influx through the plasmalemma (Silei et al., 1999; Hegg et al., 2000), hence the SERCA genes were not suppressed. Incidentally, SERCA inhibition was previously found to reduce microglial phagocytosis (Morales-Ropero et al., 2021), which is consistent with our data showing that nimodipine inhibited phagocytic activity (Fig. 4) concomitant with the downregulation of SERCA genes (Table 1).

Nimodipine consistently inhibited the phenotypic shift of microglia in activated brain slice preparations and cultures, suggesting that microglia were reliably targeted by the treatment in their neural tissue environment, in co-culture with neurons and astrocytes, and in monoculture (Figs. 1-4). We explored DEGs underlying these microglial morphological and functional phenotypic changes and provide a novel, comprehensive overview of nimodipine action at the level of gene expression in activated primary microglial cultures (Fig. 6, Table 2). Nimodipine is assumed to have achieved its effects on gene expression primarily by modulating intracellular Ca^2+^ signaling (Barbado et al., 2009), taken that Ca^2+^ is a well-known second messenger in the control of gene expression. At least three Ca^2+^-dependent gene transcriptional pathways regulated by VGCCs have been previously identified, including (i) a calmodulin-dependent kinase and cAMP response element binding protein (CREB)-dependent pathway, (ii) a pathway initiated by Ca^2+^-sensitive proteins, associated with the VGCC signaling complex, that activate transcription factors, and (iii) a pathway regulated by Ca^2+^-binding transcription factors (e.g., DREAM) (Barbado et al., 2009). In addition, a proteolytic fragment of the intracellular carboxyl terminus of the α subunit of LVGCCs (specifically of Cav1.2 and Cav 1.3 found here on microglia) can translocate to the cell nucleus to act as a transcription factor itself (Gomez-Ospina et al., 2006; Lu et al., 2015). As evidenced by our functional enrichment analysis (Fig. 7B), nimodipine is thought to finely modulate intracellular signal transduction pathways (e.g., expression of protein kinases), by inhibiting LVGCC opening to cause the gene expression changes described here.

In conclusion, this study has provided evidence that nimodipine effectively suppresses microglial activation by modulating Ca^2+^-dependent intracellular signaling cascades and gene expression involved in microglial Ca^2+^ homeostasis and immune responses. On the basis of these novel findings, consideration should be given to expanding the medical field of indication of nimodipine.

## List of abbreviations

aCSF: artificial cerebrospinal fluid
AMPA: α-amino-3-hydroxy-5-methyl-4-isoxazole-proprionic acid
ANOVA: one-way analysis of variance
BP: biological processes
DEG: differentially expressed gene
ER: endoplasmic reticulum
GO: gene ontology
Iba1: ionized calcium binding adaptor molecule 1
IFNγ: interferon-γ
IL-10: interleukin-10
IL-1β: interleukin-1β
IP3R: inositol triphosphate receptor
LPS: lipopolysaccharide
LVGCC: L-type voltage gated calcium channel
NMDA: N-methyl-D-aspartate
mOGD: mild oxygen-glucose deprivation
PCA: principal component analysis
PGE2: prostaglandin E2
Ryr: ryanoid receptor
SD: spreading depolarization
SERCA: sarcoplasmic/endoplasmic reticulum Ca^2+^-ATPase
TI: transformation index
TLR4: toll-like receptor 4
TNF-α: tumor necrosis factor-α
TRP: transient receptor potential
VGCC: voltage gated calcium channel

## Funding

The author(s) disclosed receipt of the following financial support for the research: The EU’s Horizon 2020 research and innovation program grant number 739593; the National Research, Development and Innovation Office of Hungary (grant numbers K134334, K146725, and 2023-1.1.1-PIACI_FÓKUSZ-2024-00029); the Ministry of Innovation and Technology of Hungary and the National Research, Development and Innovation Fund (grant number TKP2021-EGA-28 financed under the TKP2021-EGA funding scheme); the Ministry of Culture and Innovation of Hungary (grant number EKÖP-24-4-SZTE-622); the National Brain Research Program 3.0 of the Hungarian Academy of Sciences; and the Research Fund of the Albert Szent-Györgyi Medical School, University of Szeged, Hungary. Project no. RRF-2.3.1-21-2022-00015 has been implemented with the support provided by the European Union.

## Competing interests

The authors declare that they have no competing interests.

## Authors’ contributions

I. P.: Formal Analysis, Investigation, Visualization, Writing – original draft; V.V.: Formal Analysis, Visualization; E.Q.: Formal Analysis, Visualization, Writing – review & editing; R.F.: Formal Analysis, Investigation, Visualization; D. K.: Formal Analysis; Investigation; K.V.: Formal Analysis, Visualization; P.A.S.: Formal Analysis; Á. M.: Methodology, Supervision, Writing – review & editing; K.G.: Supervision, Writing – review & editing; F. B.: Supervision, Writing – review & editing, Funding acquisition; E. F.: Conceptualization, Formal Analysis, Visualization, Supervision, Writing – original draft, Funding acquisition.

## SUPPLEMENTARY MATERIAL

**Table S1.** Primary and secondary antibodies used for immunocytochemistry and Western blotting.

| Method | Model | Primary antibody | Company | Final dilution | Secondary antibody | Company | Final dilution |
| --- | --- | --- | --- | --- | --- | --- | --- |
| Immuno-cytochemistry | Cell culture | Rabbit anti-Iba1 polyclonal | Abcam, Cambridge, England | 1:1000 | Alexa Fluor 568 goat anti-rabbit | Invitrogen, Carlsbad, CA, USA | 1:1000 |
|  | Brain slice preparation | Rabbit anti-Iba1 polyclonal | Abcam, Cambridge, England | 1:300 | Alexa Fluor™ 488 goat anti-rabbit | Thermo Fisher Scientific, USA | 1:1000 |
| Western blot | Cell culture | Rabbit anti-Iba1 polyclonal | Abcam, Cambridge, England | 1:1000 | Anti-rabbit IgG, peroxidase conjugated | Sigma, St. Louis, MO, USA | 1:1000 |
|  |  | Mouse anti-GAPDH monoclonal | Sigma, St. Louis, USA | 1:20000 | Anti-mouse IgG, peroxidase conjug. | Sigma, St. Louis, MO, USA | 1:1000 |

**Figure S1.**
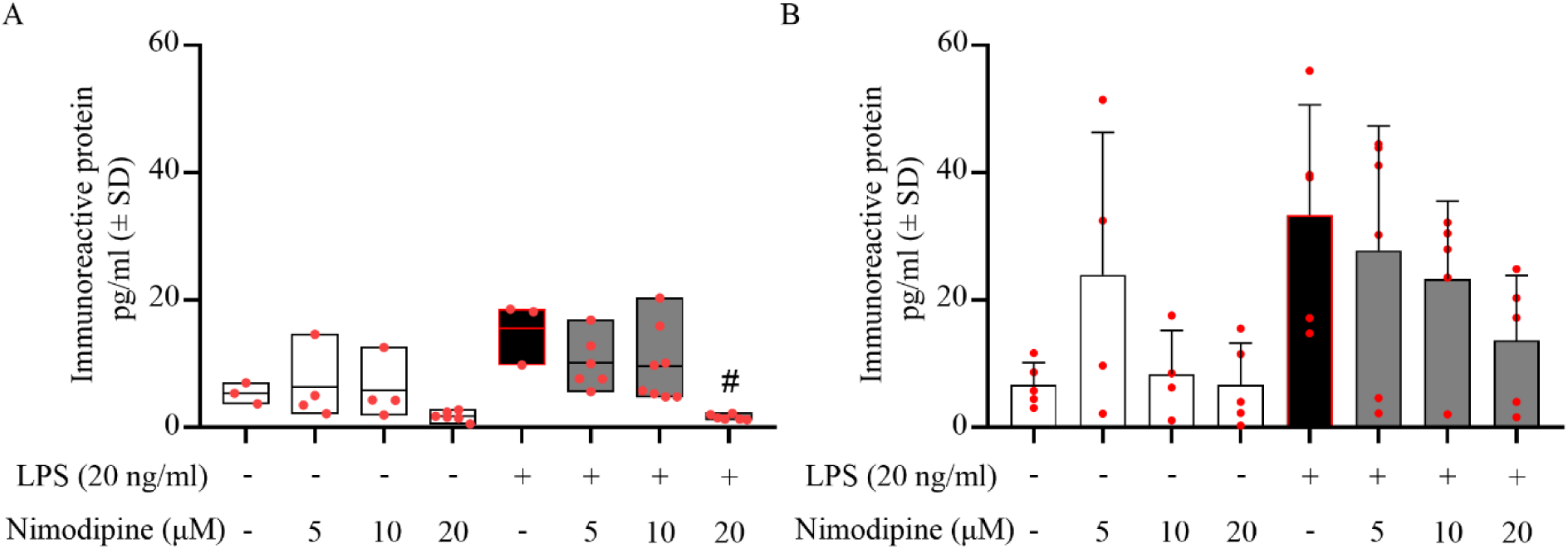
The effect of nimodipine on microglial tumor necrosis factor-α secretion (TNF-α). **A,** Quantitative analysis of TNF-α concentration in the cell culture medium of co-cultures by ELISA. **B,** Quantitative analysis of TNF-α levels in the monoculture medium by ELISA. Data are presented as mean±SD; red spheres represent individual values in each group. Normality of data distribution was determined by Shapiro-Wilk test (A, p=0.025; B, p=0.175). Data were analyzed by Kruskal-Wallis test (p<0.0002) followed by Dunn’s multiple comparison (p<0.05^#^ vs. LPS alone) (A) or one-way analysis of variance (ANOVA) (f=2.783, p=0.0229*) followed by Tukey’s multiple comparison (B).

**Figure S2.**
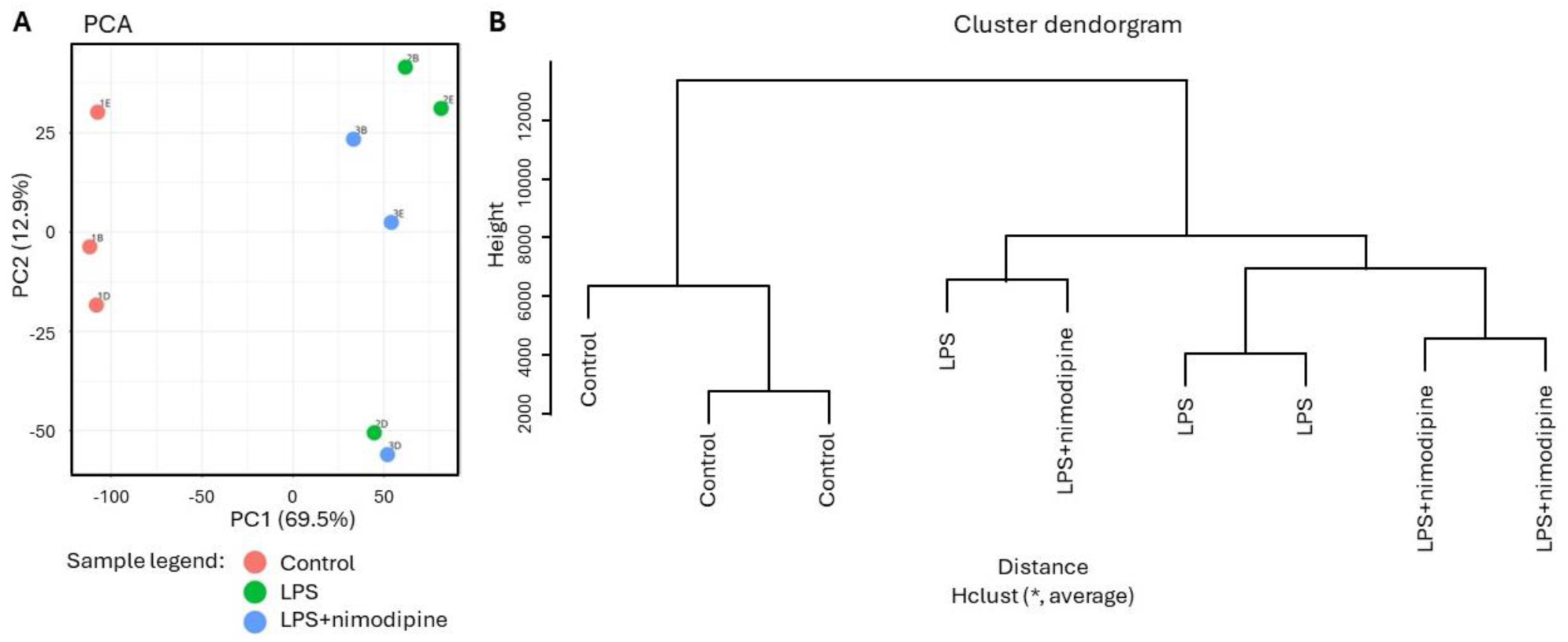
Principal component analysis (PCA) (A) and hierarchical clustering of the three replicate samples in each of the three experimental conditions.

**Figure S3.**
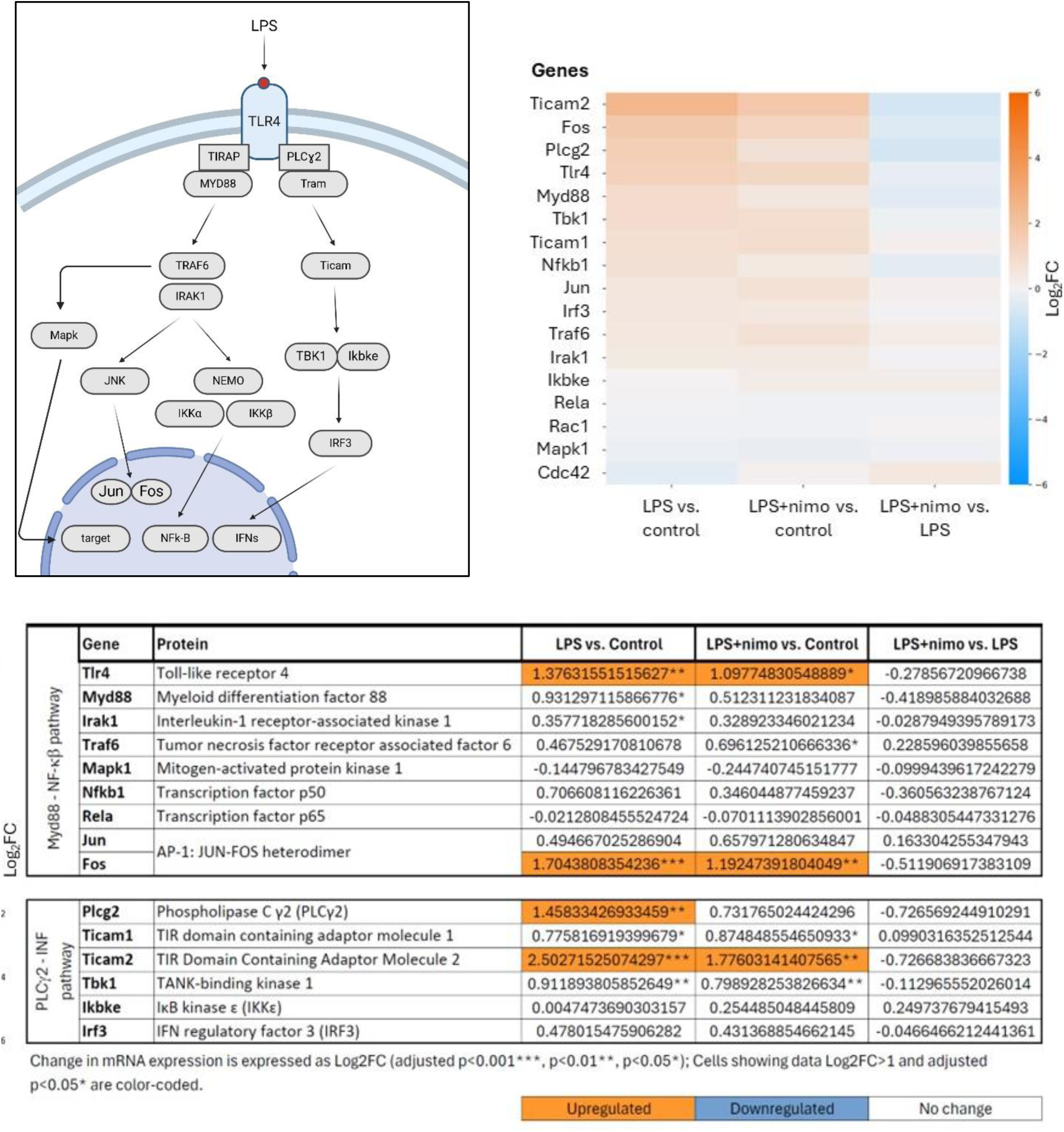
Graphical illustration, heat map and table of Log_2_FC values of differentially expressed genes (DEGs) of the TLR4 intracellular signaling cascades. The illustration was created in Biorender.

**Figure S4.**
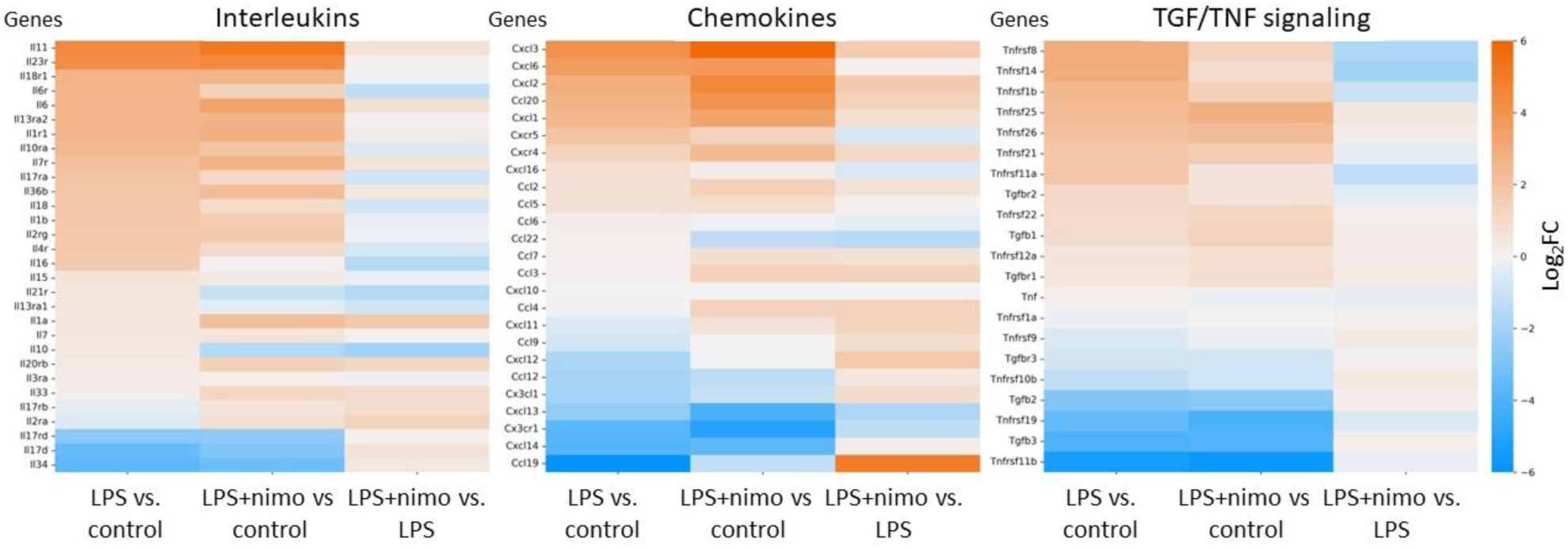
Heat maps of differentially expressed genes (DEGs) of cytokine production and cytokine receptors.

**Figure S5.**
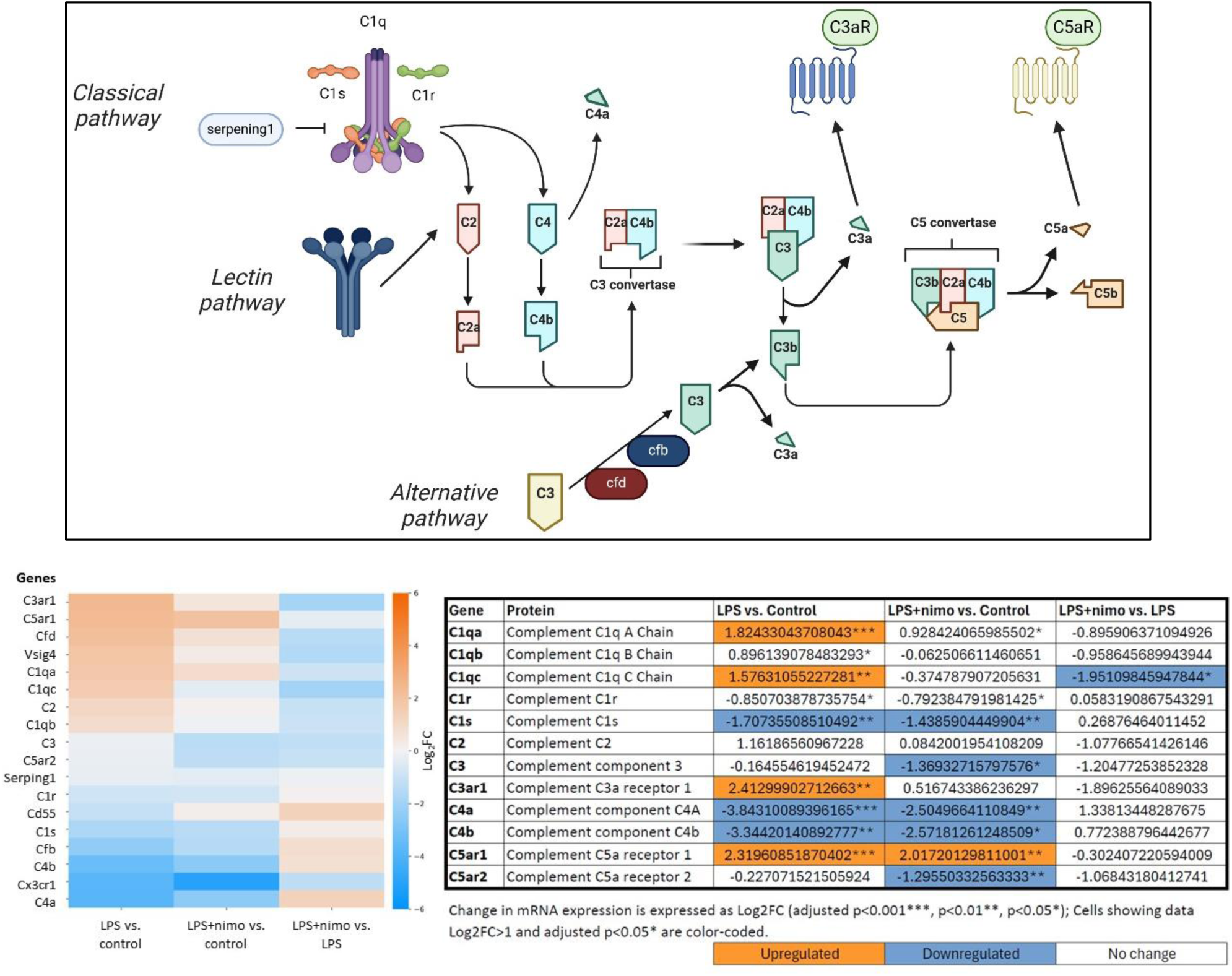
Graphical illustration of the complement cascade, and heat map and table of Log_2_FC values of differentially expressed genes (DEGs) of the complement cascade. The table represents elements of the classical pathway. The illustration was created in Biorender.

**Figure S6.**
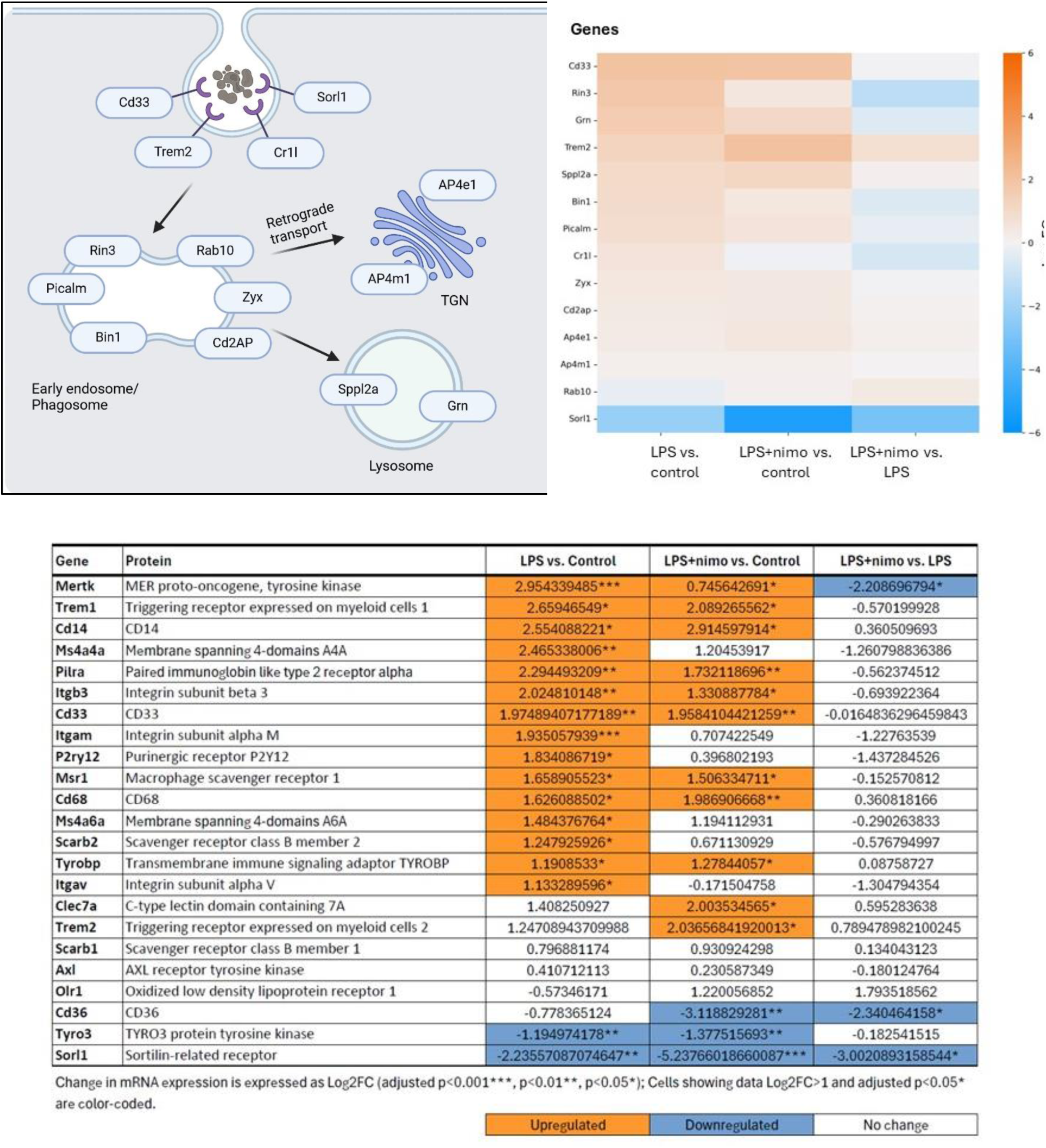
Graphical illustration, heat map and table of Log_2_FC values of differentially expressed genes (DEGs) of markers of phagocytosis. The illustration was created in Biorender.

**Figure S7.**
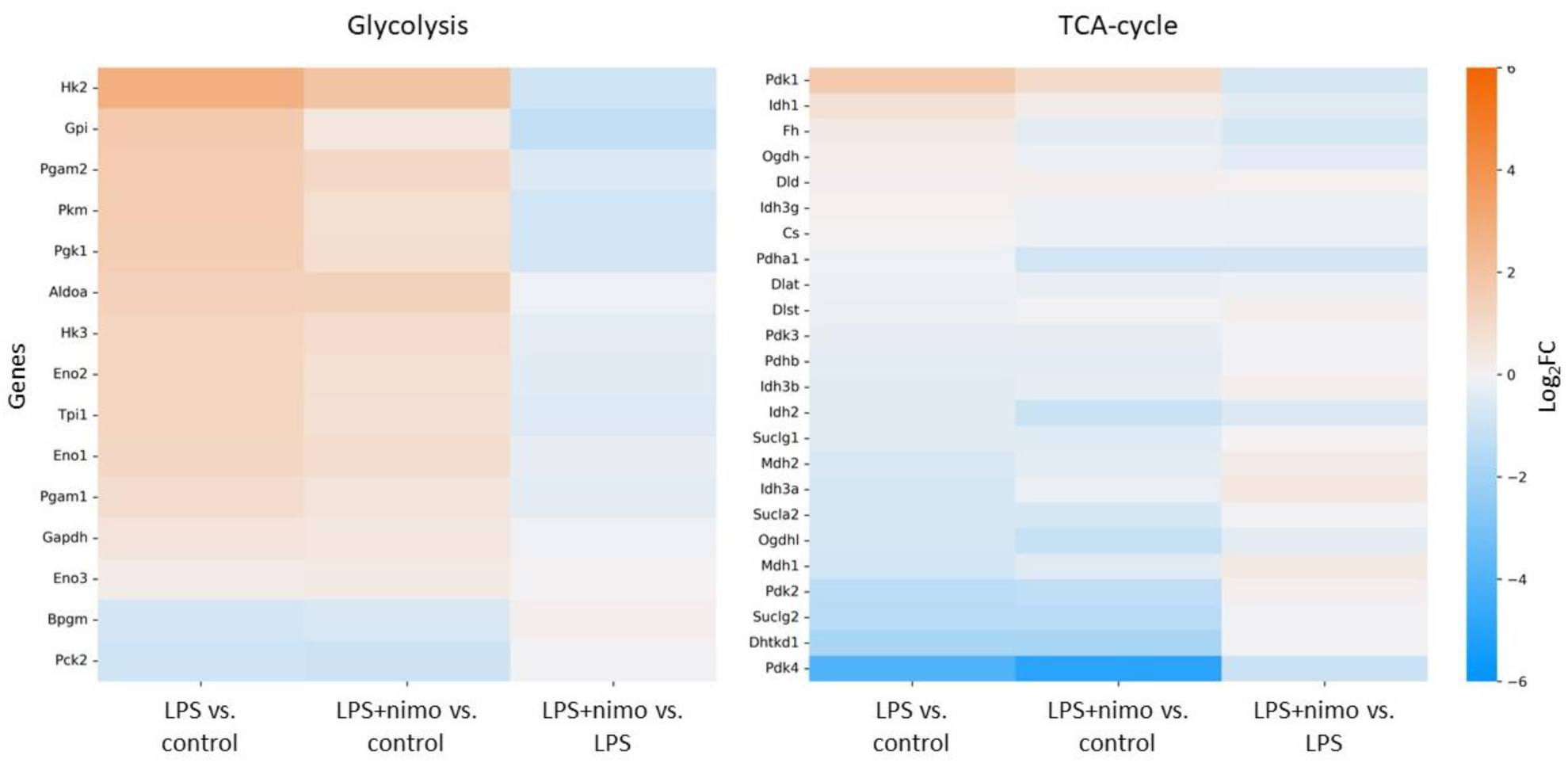
Heat map of differentially expressed genes (DEGs) of glycolysis and the mitochondrial tricarboxylic acid (TCA) cycle.

**Figure S8.**
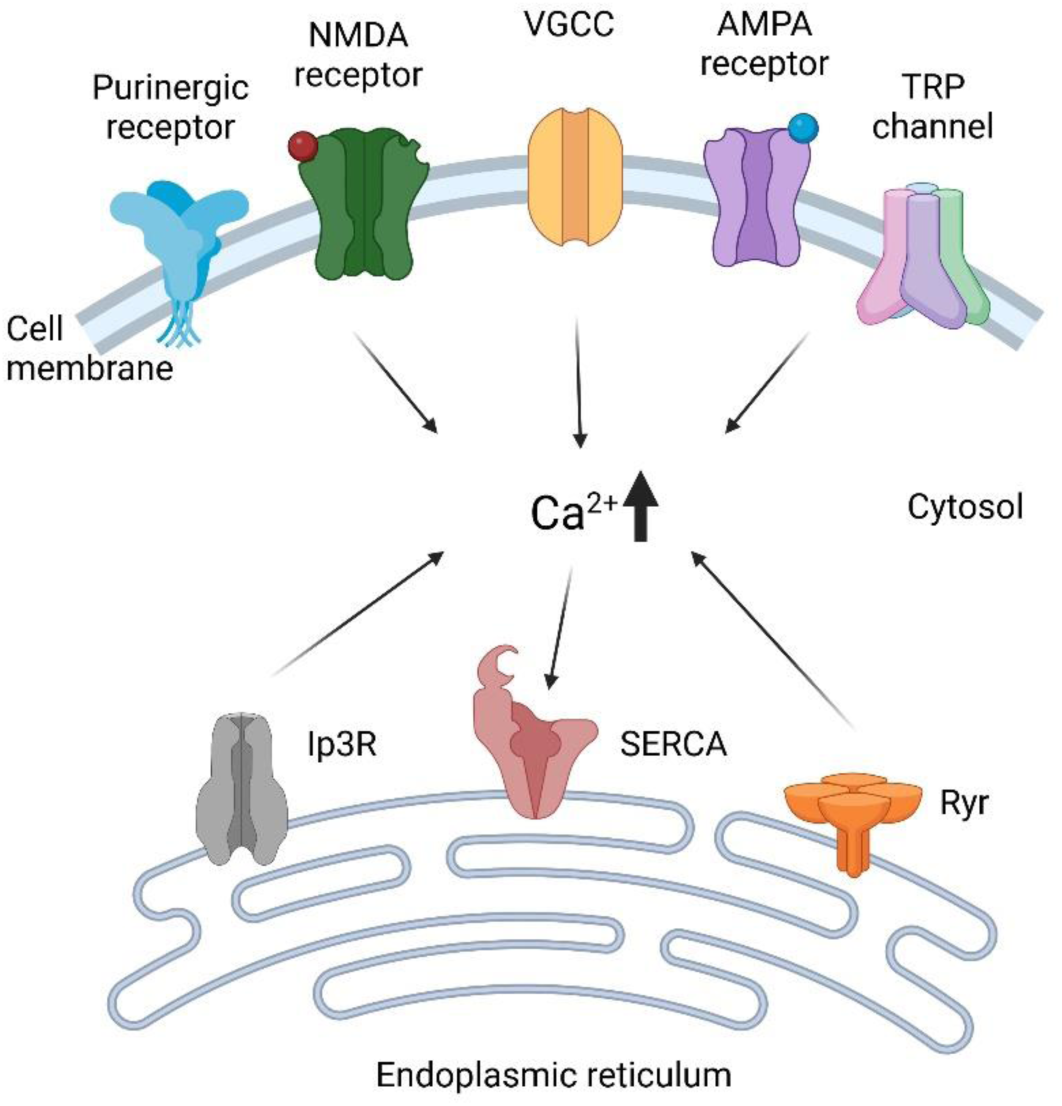
Molecular pathways of cytosolic Ca^2+^ accumulation and Ca^2+^ removal in microglia. The illustration was created in Biorender

## Notes

### Competing Interest Statement

The authors have declared no competing interest.

